# Plant Trans-Golgi Network/Early Endosome pH regulation requires Cation Chloride Cotransporter (CCC1)

**DOI:** 10.1101/2020.01.02.893073

**Authors:** Daniel W McKay, Yue Qu, Heather E McFarlane, Apriadi Situmorang, Matthew Gilliham, Stefanie Wege

## Abstract

Plant cells maintain a low luminal pH in the Trans-Golgi-Network/Early Endosome (TGN/EE), the organelle in which the secretory and endocytic pathways intersect. Impaired TGN/EE pH regulation translates into severe plant growth defects. The identity of the proton pump and proton/ion antiporters that regulate TGN/EE pH have been determined, but an essential component required to complete the TGN/EE membrane transport circuit remains unidentified − a pathway for cation and anion efflux. Here, we have used complementation, genetically encoded fluorescent sensors, and pharmacological treatments to demonstrate that the TGN/EE localised Arabidopsis Cation Chloride Cotransporter (CCC1) is this missing component necessary for regulating TGN/EE pH and function. Loss of CCC1 function leads to alterations in TGN/EE-mediated processes including endo- and exocytosis, and trafficking to the vacuole, and response to abiotic stress, consistent with the multitude of phenotypes observed in *ccc1* knockout plants. This discovery places CCC1 as a central component of plant cellular function.

## Introduction

The plant Trans-Golgi Network/Early Endosome (TGN/EE) has a complex cellular role. One of its key roles is sorting and delivering proteins to the apoplast, plasma membrane (PM) and vacuole (Dettmer et al., 2006; Viotti et al., 2010; Sze and Chanroj, 2018). This cellular function of the TGN/EE requires a finely tuned luminal pH (Luo et al., 2015; Reguera et al., 2015). Within the endomembrane system, the TGN/EE has the most acidic luminal pH, which is established by the TGN/EE-localised V-H^+^-ATPase (Martinière et al., 2013, Luo et al., 2015). Plants with reduced V-H^+^-ATPase activity (called *det3*) exhibit severe developmental growth defects, salt sensitivity, and increased TGN/EE luminal pH, which is accompanied by alterations in exocytosis of PM proteins, such as BRI1, and defects in cell division and expansion (Schumacher et al., 1999; Dettmer et al., 2006; Brüx et al., 2008; Krebs et al., 2010; Luo et al., 2015).

In addition to the V-H^+^-ATPase, endomembrane luminal pH regulation is also dependent on the activity of cation/proton exchangers (antiporters), which provide a pathway for electroneutral proton efflux from, and cation import into, the TGN/EE lumen. The best characterised cation/proton exchangers in the TGN/EE are NHX5 and NHX6. Double *nhx5/nhx6* mutants have similar, but not identical, growth defects to *det3*, with reduced cell length, a large decrease to overall plant size and increased salt sensitivity of root growth and seed germination (Bassil et al., 2011; Reguera et al., 2015). However, in contrast to *det3*, the *nhx5/nhx6* TGN/EE lumen is hyperacidic, consistent with the proposed role of NHX in exporting protons out of the TGN/EE (Luo et al., 2015; Reguera et al., 2015). As in *det3*, trafficking of BRI1 is also altered in *nhx5/nhx6*; however, only recycling is affected while secretion to the PM is not (Dragwidge et al., 2019). Similarly, the PM abundance of the membrane integrated proteins, PIN1 and PIN2, are reduced in *nhx5/nhx6* (Dragwidge et al., 2018). Moreover, the hyperacidity of *nhx5/nhx6* TGN/EE lumen results in mis-sorting of vacuolar proteins due to altered binding of targets by the Vacuolar Sorting Receptor VSR1;1 (Reguera et al., 2015).

In animals, chloride/proton antiporters of the CLC (Chloride Channel) protein family are important for lysosome acidification. This function is reliant on the coupled export of protons and import of anions, as demonstrated through the use of a CLC variant that conducted uncoupled proton/anion transport that was unable to complement the increased lysosomal pH of a mouse *clc-5* knockout (Novarino et al., 2010). In plants, two members of the CLC family, CLCd and CLCf, are localised in the Golgi and TGN/EE (Marmagne et al., 2007; von der Fecht-Bartenbach et al., 2007). CLCd is important for pathogen resistance, with otherwise minimal developmental growth defects in the single *clcd* knockout (Guo et al., 2014). It has been suggested that CLCd and CLCf fulfil similar, partially redundant, roles to their animal homologues (Sze and Chanroj, 2018).

Collectively, these observations demonstrate that a typical consequence of alterations in TGN/EE lumen pH regulation are defects in protein trafficking, particularly in protein exocytosis, changes in salt tolerance and defects in cell elongation and division, resulting in severe impacts on plant growth. The current model of TGN/EE pH regulation, however, is incomplete; it contains a proton pump (V-H^+^-ATPase) and anion- and cation-proton exchangers (CLC, NHX), but no transport protein that mediates either cation or anion efflux has yet been identified. Here, we have explored a lead that suggests CCC1 is localised to the endomembrane system, a result provided through heterologous expression studies (Colmenero-Flores et al., 2007; Henderson et al., 2015). Plant CCCs mediate electroneutral cation and anion symport, and are therefore excellent candidates to provide an ion efflux mechanism (Colmenero-Flores et al., 2007, Henderson et al., 2018). Arabidopsis contains only a single *CCC* gene, *CCC1*, whose knockout exhibits a complex and severe phenotype including large reductions in overall plant size, a bushy appearance characterised by increased axillary shoot outgrowth, frequent stem necrosis, very low fertility, alterations in pathogen response, changes in cell wall composition, and changes in seed ion concentrations (Johnson et al., 2004; Colmenero‐Flores et al., 2007; McDowell et al., 2013; Henderson et al., 2018; Han et al., 2020). In addition, *ccc* knockouts in both Arabidopsis and rice also display changes in ion distribution. Plate grown *ccc1* do not show a stronger reduction in primary root growth compared to wildtype plants under saline conditions; however, hydroponically grown Arabidopsis *ccc1* plants accumulate more Cl^−^ in shoots under both control and 50 mM NaCl conditions while soil grown *ccc1* plants accumulate more Cl^−^ in shoots when exposed to 50 mM Cl^−^ salts (Colmenero‐Flores et al., 2007; Henderson et al., 2015). Roots of *Osccc1.1* plants also had altered ion accumulation profiles under both control and saline conditions (Chen et al., 2016). Given the endomembrane localisation of Arabidopsis CCC1, the alterations in ion distribution may be the result of mechanisms similar to those that result in altered response to salinity in *det3* and *nhx5/6*.

Here we confirm the endomembrane localisation of CCC1 by stable expression in Arabidopsis, and show it is localised in the TGN/EE. We demonstrate that the non-proton coupled ion transporter, CCC1, is required for the regulation of the TGN/EE luminal pH, for the efflux of ions, and that loss of CCC1 function leads to defects in TGN/EE-dependent processes. Specifically, *ccc1* knockouts exhibit reduced rates of endomembrane trafficking. We also investigate the role of CCC1 in response to salt and osmotic stress and find that several processes in *ccc1* plants, such as germination and root hair elongation, are either more or tolerant to salt and osmotic stress or are complemented. We further find that in wildtype plants the luminal pH of the TGN/EE increases in response to osmotic shock; this does not occur in plants lacking CCC1. Overall, these results establish a link between TGN/EE pH regulation and response to osmotic and salt stress and we propose that CCC1 is a missing core component of the ion- and pH regulating machinery of the TGN/EE

## Results

### CCC1 *is ubiquitously expressed*

Previous reports on *CCC1* expression are contradictory. Promoter-GUS studies indicated that *CCC1* expression is restricted to specific tissues, such as root stele or hydathodes and pollen; while RNA transcriptomic studies, including single-cell RNAseq, suggest expression occurs in a broader range of cell types (Colmenero‐Flores et al., 2007, Wendrich et al., 2020). To clarify the tissue expression pattern of *CCC1*, we transformed wildtype Arabidopsis plants with a 2kb genomic DNA sequence upstream of the *CCC1* coding region driving the expression of nuclear-localised triple Venus (a bright variant of the yellow fluorescent protein) or β-glucuronidase (GUS), named *CCC1prom::Venus* and *CCC1prom::GUS*, respectively. Combined analysis of fluorescence and GUS-staining revealed that *CCC1* is expressed in all cell types, including all root cells, hypocotyl, leaf and stem epidermis, guard cells and trichomes, as well as mesophyll cells and all flower parts, with a particularly strong signal in stamen filaments (Fig. 1). *CCC1* promoter activity reported by Venus fluorescence, or by GUS-activity, was slightly different despite use of the identical promoter sequence. For instance, fluorescence was detectable in root cortex and epidermis cells, including root hairs, and in the gynoecium, while GUS staining did not indicate expression in these cells. This is likely due to the increased sensitivity of the Venus fluorescence method.

**Figure 1.**
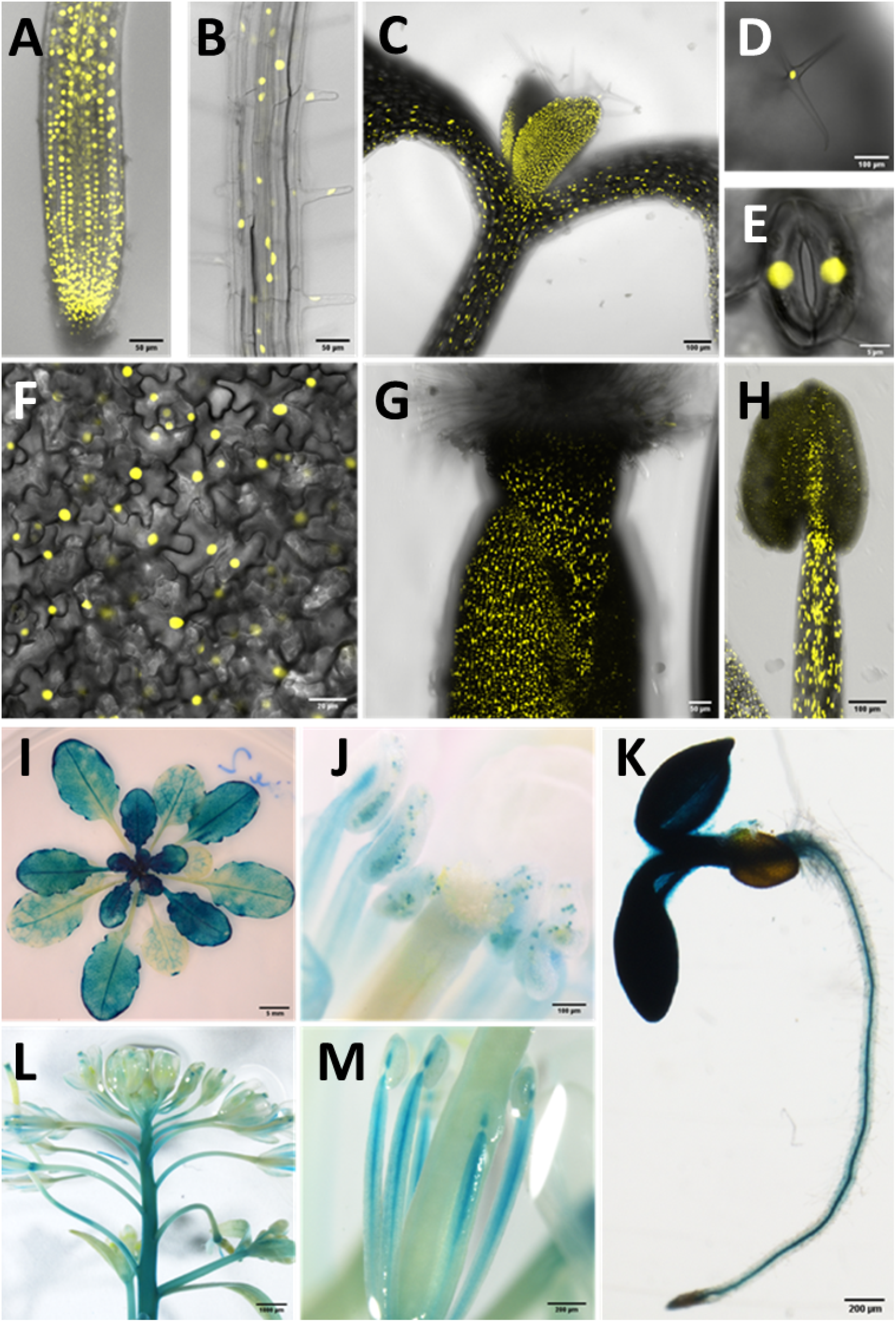
*CCC1* is expressed in most cell types. *CCC1* promoter driven expression of either NLS-Venus (bright YFP variant with a nuclear localisation signal) or β-glucuronidase (blue GUS staining). A-H) NLS-Venus expression indicating *CCC1* promoter activity in all root cells, including (A) the root tip, (B) root epidermal cells; (C) hypocotyl; all leaf cells including (D) trichomes, (E) guard cells, and (F) leaf epidermal and mesophyll cells; and reproductive organs (G) gynoecium and (H) stamen tissues. I-M) GUS staining indicating promoter activity predominantly in (I) younger leaves, (J) anthers, (K) root stele, (L) floral stem, and (M) stamen. Scale bars are 50 μm (images A, B, G), 100 μm (images C, D, H, J), 5 μm (image E), 20 μm (image F), 5 mm (image I), 200 μm (images K, M) and 1000 μm (image L).

### Loss of CCC1 results in growth defects in root cells

*ccc1* knockout plants are severely affected in their growth, including a reduced shoot size and shorter primary roots (Fig. 2A, and previously shown by Colmenero‐Flores et al., 2007; Henderson et al., 2015). We investigated the origin of the root phenotype of *ccc1* at a cellular level and found that CCC1 function is required for cell elongation. Knockouts develop both shorter root epidermis cells, and shorter root hairs (Fig. 2B-E). *ccc1* knockouts also show a complete lack of collet hairs (Fig. 2F), which are epidermal root hairs formed in some plant species in the transition zone between the root and the hypocotyl (Sliwinska et al., 2015). In addition, *ccc1* root hairs also displayed branching and bulging, although, at a low frequency (Fig. S1). Ruptured root hairs were not observed. Independent of the defect of root hair elongation, *ccc1* plants frequently developed root hairs in cell files that usually exclusively contain atrichoblasts (non-root hair cells). The trichoblast marker *PRP3::H2B-2×mCherry* was used to confirm root hair cell identify and showed the frequent presence of multiple adjacent trichoblasts in *ccc1*, which was absent from wildtype plants (Fig. S1).

**Figure 2.**
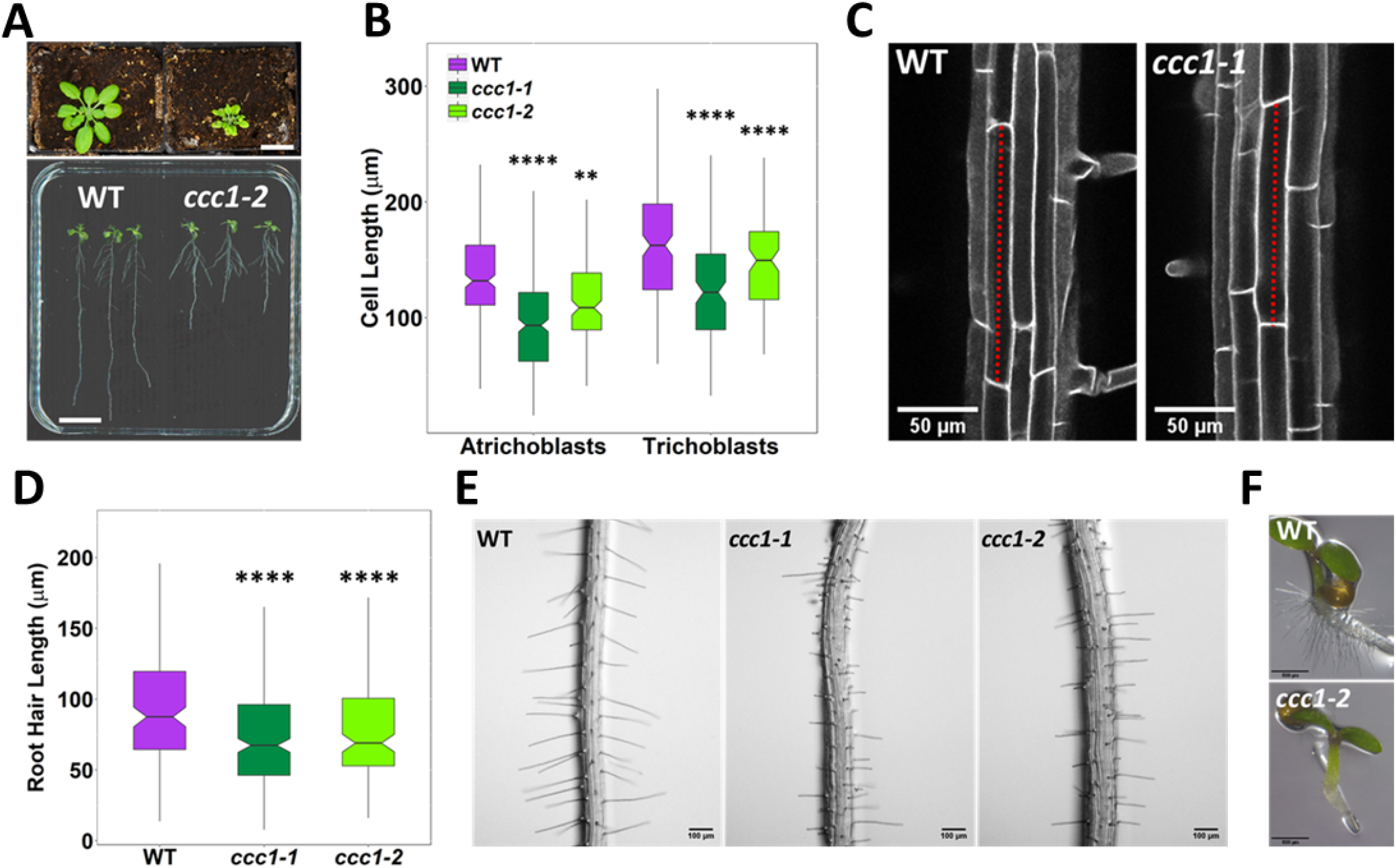
*ccc1* plants show defects in cell elongation. A) Top image, *ccc1* (right) have smaller shoots and deformed leaves compared to wildtype (le ft) plants. Plants grown 26 d in short day, scale bar is 2 cm. Bottom image, *ccc1* (right) have shorter primary roots compared to wildtype (left) plants. Plants grown 14 d in long day, scale bar is 2 cm. B-C) Root epidermal cells are shorter in 2 *ccc1* lines. n > 13 plants. Images are maximum intensity projections of cell wall autofluorescence. Scale bars are 50 μm.-D-E) *ccc1* plants have shorter root hairs. n > 900 root hairs of > 30 plants. Scale bars are 100 μm. F) *ccc1* pla nts do not develop collet hairs. Scale bars are 500 μm. Boxplots show range excluding outliers; median, 1st and 3rd quartile are indicated. Points represent individual measurements. Student’s t-tests comparing *ccc1* to wildtype. ** indicates *P* < 0.01, **** indicates *P* < 0.0001.

### Functional GFP-CCC1 localises to the endomembrane system

We had previously localised CCC1-GFP to the Golgi and TGN/EE in transient expression assays in *Nicotiana benthamiana* (Henderson et al., 2015). In contrast, other studies have suggested that CCC1 might localise to the PM, which has led to multiple and conflicting interpretations of CCC1 function (Wegner, 2017; Domingos et al., 2019). To clarify the subcellular localisation of CCC1, we generated plants that stably express N-terminally tagged GFP-CCC1, using the *EXP7* (*Expansin7*) trichoblast-specific promoter to express GFP-CCC1 in root epidermal cells. This approach was adopted after many attempts to generate plants with native *CCC1* promoter driven CCC1 expression, which did not produce any transformants. Approaches included the use of different protein linkers, different fluorescent proteins, smaller tags such as FLAG-tag, and different tag locations (N- and C-terminal, and internal). The difficulty in obtaining transformed plants might suggest that tagging interferes with CCC1 function in embryonic or meristematic tissue where it is highly expressed (Fig. 1); while internal tagging might have disrupted protein folding as transformants were recovered, but no fluorescence could be detected. We therefore decided to express GFP-CCC1 in a mature cell type, in which we had identified a clear phenotypic defect in *ccc1* − trichoblasts. Expression in these epidermal cells was successful and importantly, complemented the short root hair phenotype of *ccc1* knockout plants (Fig. 3).

**Figure 3.**
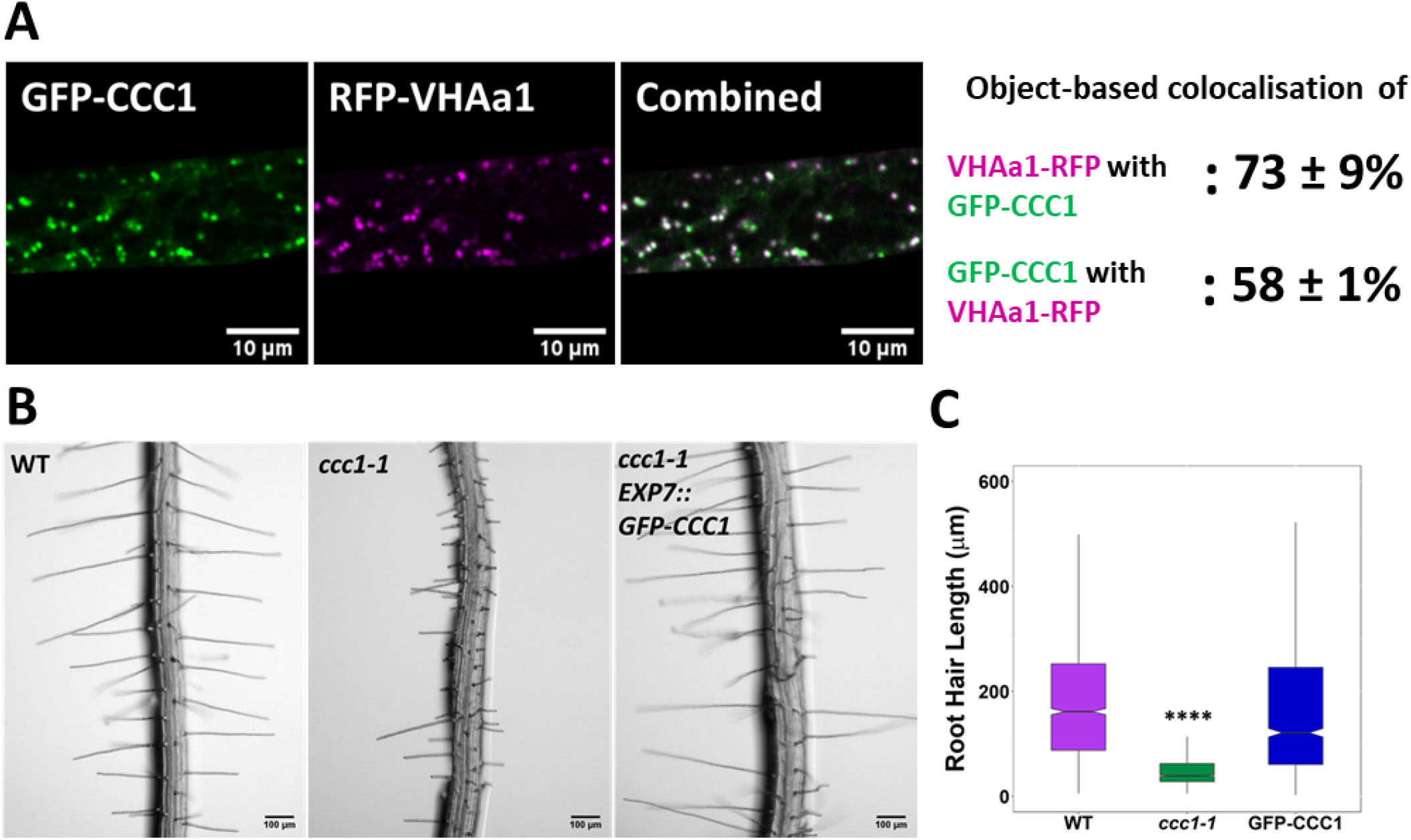
Stably expressed GFP-CCC1 is functiona l and localised in the TGN/EE. A) GFP-CCC1 (green) and VHAal-RFP (magenta) colocalise. Colocalisation was calculated using DiAna object-based colocalisation plugin in lmageJ; Pearson’s coefficient was also calculated as 0.86 ± 0.055. Error is standard deviation. n = 15 cells of 5 plants. Scale bars are 10 μm. Images are single representative optical sections from a stack. B-C) Expression of GFP-CCC1 rescues *ccc1* root hair length defects. n > 1300 root hairs. Scale bars are 100 μm. Boxplot shows range excluding outliers; median, 1st and 3rd quartile are indicated. Student’s t-tests comparing genotypes to wildtype. **** indicates *P* < 0.0001.

Stable expression of GFP-CCC1 in a native cell type revealed a similar pattern to what we previously observed in *Nicotiana benthamiana,* with the GFP signal localised to internal organelles (Henderson et al., 2015). Time-lapse imaging of GFP-CCC1 movement was consistent with what could be expected for the Golgi or TGN/EE, however, GFP-CCC1 labelled organelles did not resemble the Golgi (Videos V1 and V2). To identify the observed GFP-CCC1 labelled compartments, we crossed the stably expressed marker, VHAa1-RFP, into plants expressing GFP-CCC1 (Fig. 3A). Colocalisation of GFP-CCC1 and VHAa1-RFP was measured using object-based colocalisation analysis, with the ImageJ plugin DiAna, which revealed that 73±9% of VHAa1-RFP colocalised with GFP-CCC1, while 58±11% of GFP-CCC1 colocalised with VHAa1-RFP. The asymmetrical colocalisation indicates that, in addition to the TGN/EE, CCC1 might also localise to additional organelles of the endomembrane system (Fig. 3A), similar to NHX5 and NHX6 (Reguera et al., 2015). In addition, the Pearson correlation coefficient of pixel signal intensity for RFP and GFP channels was calculated, which gave a correlation value of 0.86±0.055. Pharmacological treatment further confirmed the endosomal localisation. Treatment with the trafficking inhibitor, brefeldin-A (BFA), caused the GFP signal to accumulate in the centre of BFA bodies, consistent with a TGN/EE localisation (Fig. S2).

We then investigated if CCC1 might cycle between the TGN/EE and PM, but the signal intensity at the PM was too low to detect. Cases like this have been observed for other ion transporters that localise mainly in endosomes but function at the PM, such as the iron transporter, IRT1. PM localisation of IRT1 can be visualised using pharmacological treatment with TyrA23 (Tyrphostin A23; Barberon et al., 2011). For CCC1, no change in the subcellular localisation was observed after treatment with TyrA23, including no signal at the PM (Fig. S2). Similarly, an osmotic shock treatment, which can sometimes induce a change in protein localisation, did not lead to any observable changes in GFP-CCC1 localisation (Fig. S2). Combined, our results show compelling evidence that CCC1 functions in the TGN/EE.

Loss or knockdown of other TGN/EE-localised proteins, such as the H^+-^V-ATPase, can affect the TGN/EE morphology (Dettmer et al., 2006). High-pressure freezing, freeze substitution, and transmission electron microscopy revealed that the lack of CCC1 did not lead to obvious morphological changes in the TGN/EE ultrastructure, and the appearances of all organelles was similar between *ccc1* mutant and wildtype cells (Fig. S2). This suggests that the defects observed in *ccc1* knockouts are probably connected to changes in TGN/EE luminal conditions.

### *Hyperosmotic stress rescues root hair elongation and improves seed germination of* ccc1

Knockouts of the TGN/EE localised ion transporters, NHX5 and NHX6, as well as the proton pump V-H^+^-ATPase are salt sensitive, with germination and root growth phenotypes (Krebs et al., 2010; Bassil et al., 2011). We therefore further investigated the response of *ccc1* plants to salt stress. As rice OsCCC1.1 is implicated in maintaining cell sap osmolality, we also investigated the response to osmotic stress.

We first assayed germination of *ccc1* on media with added salt (NaCl), or an isosmotic concentration of mannitol or sorbitol, to determine if any changes in tolerance were the result of ionic or osmotic stress (Fig 4A-B and Fig S3). Wildtype and *ccc1* seeds were germinated on media containing 0 to 300 mM of NaCl or 0 to 600 mM of mannitol/sorbitol. *ccc1* seeds typically germinated slightly earlier than wildtype seeds so germination of seeds was assessed 8 d after imbibition (= 6 days after transfer to the growth chamber). Under control conditions, over 90% of wildtype, *ccc1-1* and *ccc1-2* seeds germinated. The germination rate of all genotypes decreased with higher salt, mannitol and sorbitol concentrations, yet, this decrease was smaller in the *ccc1* knockouts than wildtype. On 450 mM mannitol, 30% ± 7% (SEM) of wildtype seeds germinated, while 79% ± 6% of *ccc1-1* and 64% ± 7% of *ccc1-2* seeds germinated. On 600 mM mannitol, 4% ± 2% of the wildtype seeds germinated, while 27% ± 6% of *ccc1-1* and 20% ± 6% of *ccc1-2* seeds were able to germinate in this condition. Similar results were obtained on sorbitol (Fig S3). A similar trend was also observable when plants were grown on media containing NaCl. Growth and germination of both wildtype and *ccc1* plants on NaCl was lower than observed on isosmotic concentrations of mannitol and sorbitol. In addition, the increased tolerance of *ccc1* seed germination to NaCl was less pronounced than on mannitol and sorbitol. At 150 mM NaCl, 67% ± 7% of wildtype seeds germinated compared with 85% ± 5% of *ccc1-1* and 95% ± 2% of *ccc1-2* seeds.

**Figure 4.**
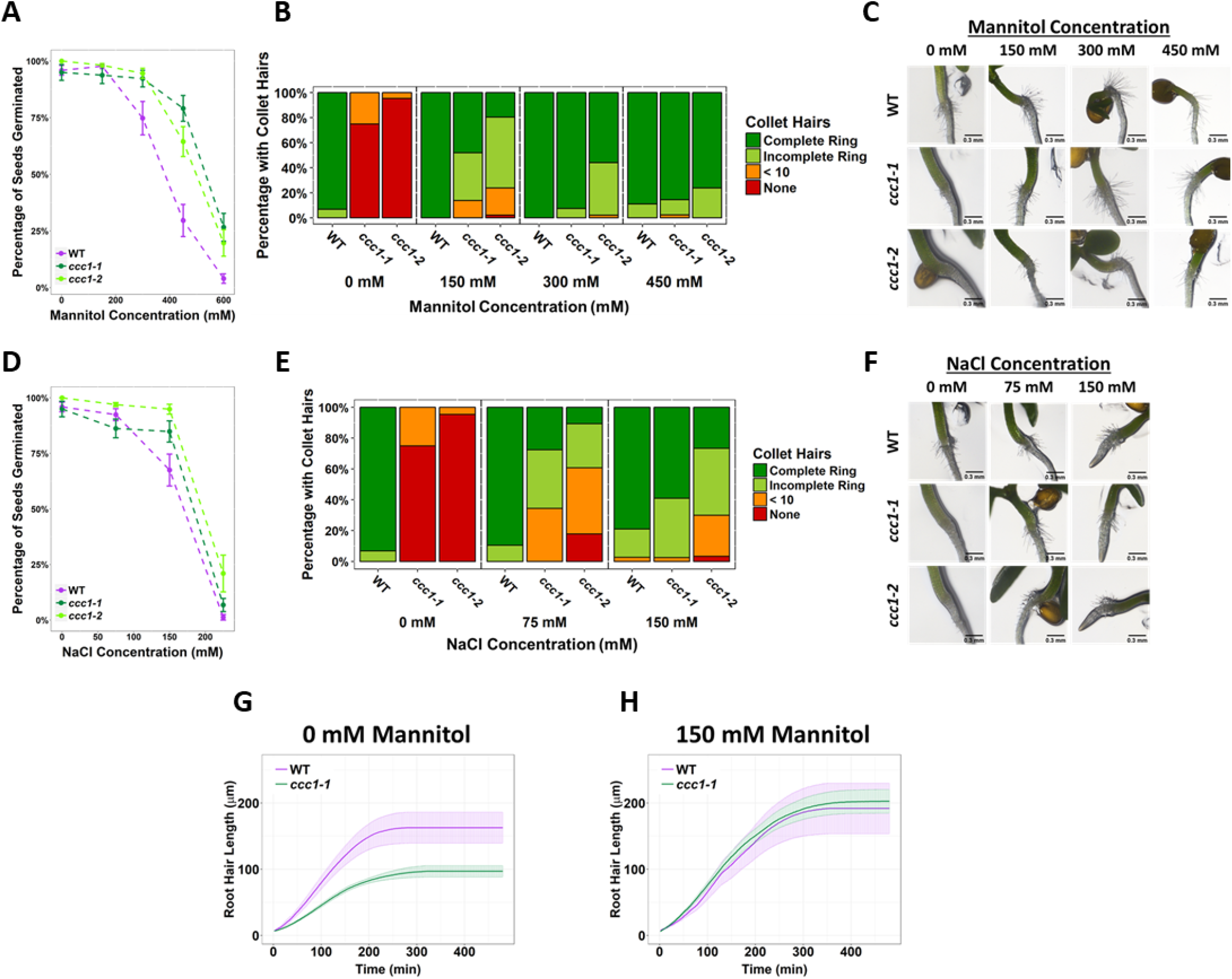
Increased external osmolality rescues cell elongation defects in *ccc1.* A, D) A higher percentage of *ccc1* seeds germinate on media with a higher osmolality, adjusted with increasing A) mannitol or D) NaCl concentrations. Number of germinated seeds assessed 6 d after imbibition. n > 90 seeds. B-C, E-F) Collet hair formation in *ccc1* is rescued on media with a higher osmolality. n > 9 plants for 450 mM treatments, > 30 plants for all other treatments. Scale bars are 300 μm. Germination and collet hair assays with mannitol, sorbitol and NaCl were performed together and therefore share the same control (0 mM). G-H) The slower rate of root hair elongation in *ccc1* is rescued when grown in media with 150 mM mannitol (See videos V3-6).

In addition to the higher germination rate of *ccc1* seeds under high osmolality, we observed that a striking phenotype of *ccc1*, the absence of collet hair formation (Fig. 2), is rescued when *ccc1* is grown on media with a greater external osmolality. Under control conditions, over 90% of wildtype seedlings had a complete ring of collet hairs while they were present in 0% of *ccc1* seedlings of either knockout allele (Fig 4C-F and Fig S3). When grown on 150 mM mannitol, 86% of *ccc1-1* seedlings and 75% of *ccc1-2* seedlings had collet hairs present as either a partial or complete ring around the hypocotyl. This improved to 100% of *ccc1-1* and 98% of *ccc1-2* on media with 300 mM mannitol. Like with germination, little difference was seen between mannitol and sorbitol treatments (Fig S3). NaCl treatment also resulted in the presence of collet hairs in *ccc1* knockouts, however, not to the same magnitude as observed in mannitol and sorbitol treatments. When treated with 150 mM NaCl, 98% of *ccc1-1* and 63% of *ccc1-2* had collet hairs.

*ccc1* mutants, as well as having no collet hair formation, also had shorter root hairs under control conditions (Fig. 2). Therefore, we investigated if the reduced root hair growth of *ccc1* plants is also rescued by increasing external osmolality. Root hairs below a final length of 50 μm were excluded, this consists of all root hairs which initiated, but did not elongate, as it was not possible to acquire an accurate measurement of elongation rate from such root hairs. Time lapse microscopy revealed that *ccc1* root hairs grow for the same duration as wildtype hairs, but grow at a reduced speed leading to a reduced final root hair length (Fig. 4G, videos V3-V6). Between 50 and 100 min after elongation initiated, wildtype root hairs had an average elongation rate of 0.88 ± 0.27 μm min^−1^, while *ccc1* root hairs elongated at half that speed, at 0.47 ± 0.08 μm min^−1^ (Fig. 4G). Strikingly, when grown in media with 150 mM mannitol, the elongation rate of *ccc1* root hairs was increased to be the same rate as wildtype. Final length of root hairs was therefore also rescued.

### ccc1 *cells do not have higher cell sap osmolality but are more resistant to plasmolysis*

We next investigated the origin of rescue of *ccc1* root hair elongation when grown on media with an elevated osmolality. Conceivably, this could be connected to an increased osmolality in *ccc1* cells. However, we found that cell sap osmolality was not higher in *ccc1* compared to wildtype. Moreover, it might be potentially lower, as one knockout allele was significantly different from wildtype, however, not the other (Fig. 5A).

**Figure 5.**
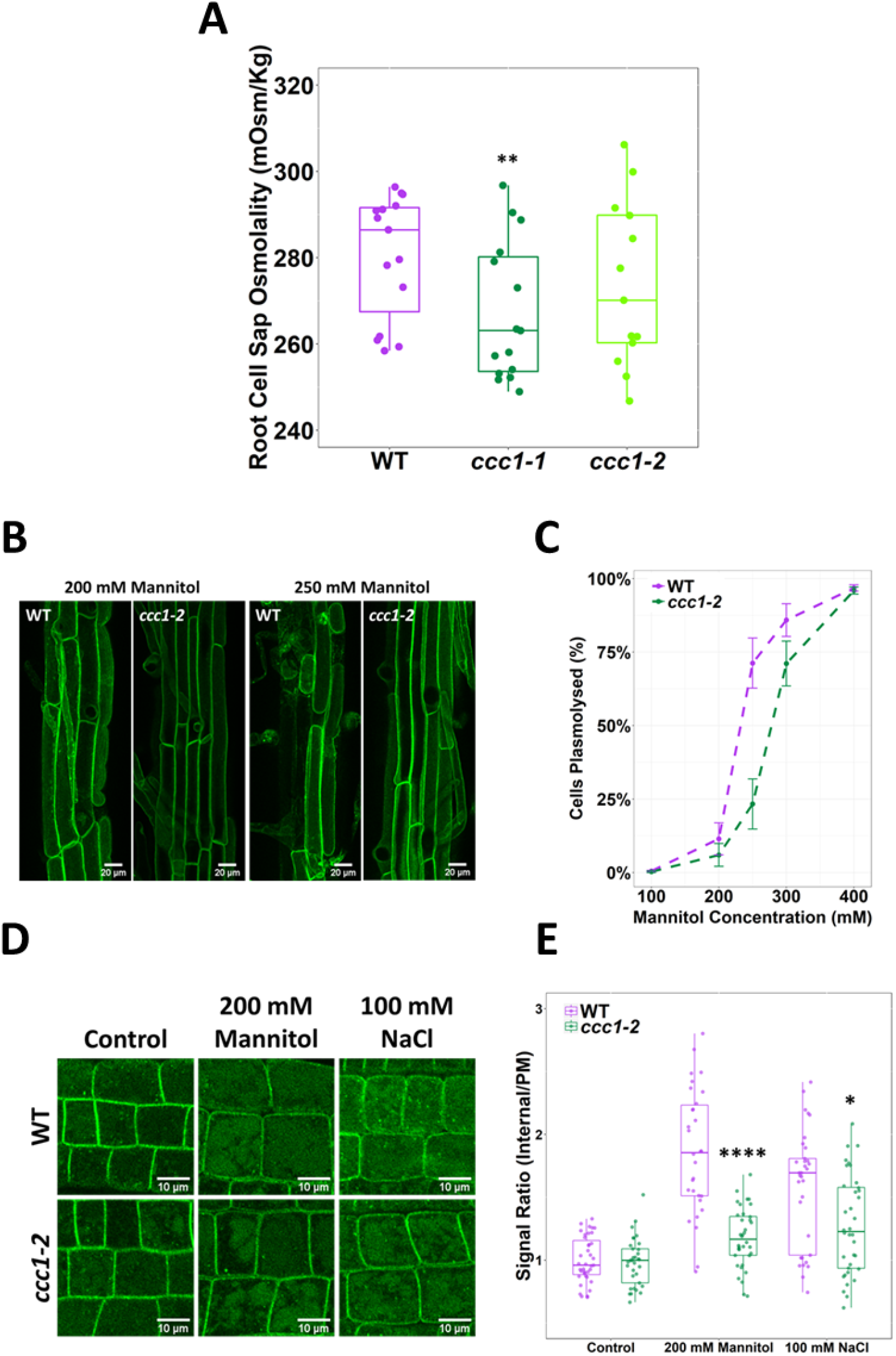
ccc1 root cells do not have a higher osmolality but are more tolerant to plasmolysis. A) The cell sap osmolality of whole root fluid, was measured using a freeze point osmomemter; no difference between wildtype and *ccc1-2* cell sap was found, while *ccc1-1* cells had a slightly lower cell sap osmolality. n = 13. B-C) *ccc1* root epidermal cells require a higher external mannitol concentration to induce incipient plasmolysis. Plasmolysed cells were counted 1 h after treatment. The PM of cells was visualised with the PM marker, GFP-LIT6b (green). 40 cells were assessed per plant, 12 plants were counted for each genotype/treatment. Scale bars are 20 μM. D-E) The internalisation of PIP2;1-GFP in response to osmotic (200 mM mannitol) and salt (100 mM NaCl) shock treatment was assayed in root epidermal cells of the elongation zone. Plants were imaged 30 min after treatment. A signal ratio of internal and PM signal was used to measure internalisation.n > 45. Scale bars are 10 μM. Boxplot shows range excluding outliers; median, 1st and 3rd quartile are indicated. Student’s t-tests comparing genotypes to wildtype. * indicates P < 0.05, ** indicates P < 0.01, **** indicates P < 0.0001

Other factors that could contribute to the improved growth of *ccc1* plants on media with high osmotic strength include altered osmoregulatory capacity. To explore if the osmoregulatory capacity of *ccc1* cells may be altered, the point of incipient plasmolysis was determined for wildtype and *ccc1* epidermal root cells. This determines the external osmotic strength required to induce plasmolysis in 50% of the cells. Plasmolysis of mature epidermal root cells was induced by submerging roots in liquid media containing high concentrations of mannitol for 30 min after which plamolysed cells were observed using plants that stably express a PM-marker. A lower proportion of plasmolysed cells were counted in *ccc1* at all but the highest mannitol concentration (Fig. 5B-C). The greatest difference was observed at 250 mM mannitol, where 71% ± 9% (SEM) of wildtype cells were plasmolysed compared to only 23% ± 9% of *ccc1* cells at the same concentration. The estimated mannitol concentration for incipient plasmolysis was determined using a fitted curve. For wildtype, 50% of cells are estimated to plasmolyse at 232 mM mannitol, while plasmolysis of 50% of *ccc1* root cells was estimated to require a mannitol concentration of 277 mM, 45 mM higher than that of the wildtype. As a higher cell sap osmolality is excluded as a reason for higher resistance to plasmolysis in *ccc1*, other factors might be the cause for the observed differences between the wildtype and *ccc1*.

### Trafficking of PIP2;1 in response to osmotic shock is reduced

In response to osmotic stress, aquaporins are removed from the PM (Boursiac et al., 2008; Hachez et al., 2014). The aquaporin PIP2;1 cycles between the PM and TGN/EE and is internalised in response to osmotic stress (Luu et al., 2012). We investigated osmotically induced internalisation of stably expressed PIP2;1-GFP in *ccc1* and wildtype plants. Wildtype and *ccc1* roots were treated with 100 mM NaCl or 200 mM mannitol for 30 min before internalisation of PIP2;1-GFP was measured by obtaining the ratio of fluorescent signal on the PM to signal inside the cell. In both wildtype and *ccc1* roots, PIP2;1-GFP remained localised to the PM when treated with only 1/2 MS (Fig. 5D-E). When treated with NaCl or mannitol, both genotypes internalised PIP2;1-GFP, but the relative portion of PIP2;1-GFP internalisation was lower in *ccc1* than wildtype, excluding a more efficient removal of PIP2;1 from the PM as the reason for the reduced plasmolysis in *ccc1* knockouts (Fig. 5D-E). Additional factors, such as the previously observed cell wall changes, might therefore contribute to observed improvement of the *ccc1* knockout when grown in media with higher external osmolality.

### Loss of CCC1 results in defects of endomembrane trafficking

The reduced internalisation of PIP2;1-GFP in *ccc1* cells in response to osmotic stress may be the result of a reduced rate of endomembrane trafficking in *ccc1* cells. We therefore investigated if *ccc1* is important for maintaining normal rates of endomembrane trafficking.

The rate of protein exocytosis, which is a combination of trafficking from secretion and recycling, was assayed using wildtype and *ccc1* lines stably expressing PIN2-GFP. BFA treatment resulted in the accumulation of PIN2-GFP in the endomembrane system, measured as a strong increase in cytoplasm:PM signal ratio. This increase was more pronounced in *ccc1* than in wildtype cells (Fig. 6A-B). BFA treatment in Arabidopsis root cells leads to the formation of BFA bodies, which are an amalgamation of Golgi and TGN/EE (Geldner et al., 2001). Interestingly, while wildtype BFA bodies were compact and bright in wildtype plants, BFA bodies in *ccc1* were less dense with a more diffuse GFP signal pattern. Upon BFA washout, wildtype plants exhibited an almost complete recovery of the PIN2-GFP cytoplasm:PM signal ratio, suggesting that protein secretion and recycling had resumed. In contrast, *ccc1* showed minimal recovery in the cytoplasm:PM signal ratio indicating a considerable decrease in PIN2-GFP exocytosis (Fig. 6A-B).

**Figure 6.**
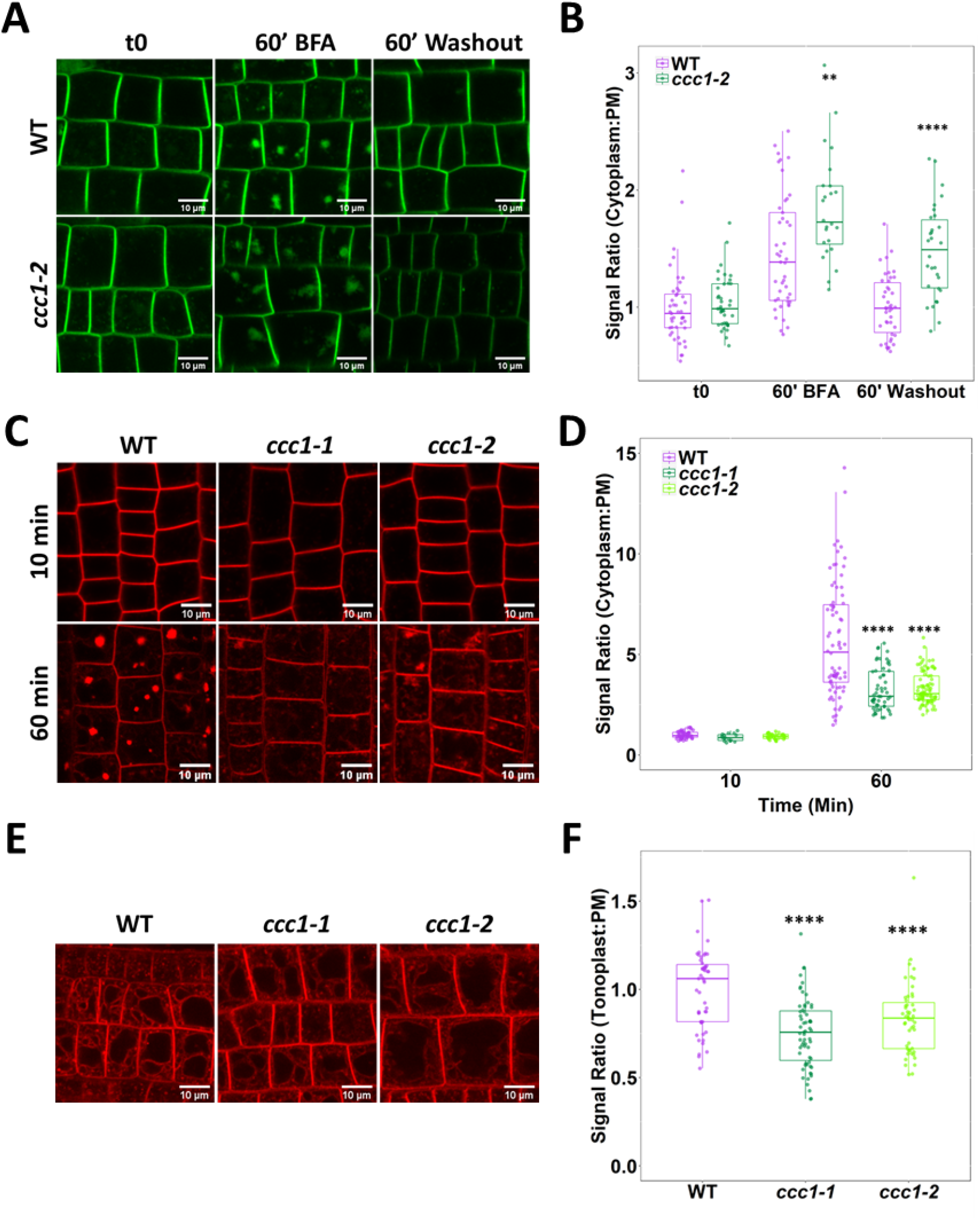
Loss of CCC1 leads to defects in endo- and exocytosis. A-B) *ccc1* root epidermal cells show a reduced recovery of PIN2-GFP (green) cytoplasm:PM signal ratio after a 60 min treatment with 25 μM BFA. n > 24 cells, 3 cells measured per plant. C-D) Endocytosis of the membrane dye FM4-64 (red), measured as an increase in the cytoplasm:PM ratio, is reduced in *ccc1* lines. Plants were kept in 25 μM BFA for the duration of the experiment. n > 56 cells of > 9 plants. E-F) Endocytosis and trafficking of FM4-64 (red) to the vacuole in the absence of BFA, measured as an increase of the tonoplast:PM signal ratio, is reduced in *ccc1* lines. n > 51 cells of > 18 plants. Boxplots show range excluding outliers; median, 1st and 3rd quartile are indicated. Points represent individual measurements. All images are single representative optical sections. Student’s t-tests comparing *ccc1* to wildtype. ** indicates *P* < 0.01, **** indicates *P* < 0.0001. All scale bars are 10 μm.

As the rate of PIP2;1-GFP internalisation was reduced, we also measured endocytosis. Endocytosis was quantified by internalisation of the endocytic tracer FM4-64, measured by an increase in the cytoplasm:PM signal ratio, both with and without BFA. After 60 min of treatment (10 min with FM4-64 & BFA, 50 min with BFA only), the fluorescence ratio was much lower in *ccc1* than wildtype cells, consistent with a large reduction in the rate of endocytosis (Fig. 6C-D). Like in the PIN2-GFP experiment, BFA bodies in *ccc1* were more diffuse than in wildtype cells. The *ccc1* PIN2-GFP BFA bodies were brighter than wildtype and the FM4-64 BFA bodies were duller than wildtype indicating that the difference in BFA body morphology in wildtype and *ccc1* was not the cause of the differences in signal internalisation in either experiment. In a second FM4-64 pulse chase experiment, trafficking of FM4-64 to the vacuole was measured without the use of BFA. Labelling of the tonoplast with FM4-64 became visible in both wildtype and mutant cells after 3 h. The reduced ratio of tonoplast:PM signal in *ccc1* compared with wildtype cells indicated a reduction in the trafficking of endocytosed FM4-64 to the vacuole (Fig. 6E-F). These observations indicate that *ccc1* cells have generic trafficking defects.

### CCC1 contributes to TGN/EE luminal pH regulation

Here, we show that GFP-CCC1 is localised to the TGN/EE and identified that CCC1 is important for endomembrane trafficking (Fig. 3 and 6). We hypothesised that the symporter might impact endomembrane trafficking through its role in regulating ion movement across the TGN/EE membrane, which may contribute to an altered luminal pH by affecting the activities of the cation- and anion-proton exchangers, and therefore ultimately the proton pump. To investigate this possible role of CCC1 in the pH regulatory network, we used both pharmacological treatments and a stably expressed TGN/EE fluorescent pH sensor.

The ionophore monensin was utilised to assess if the loss of CCC1 results in changes to the ability of the TGN/EE to maintain a stable luminal environment. Monensin is a monovalent cation ionophore, acting as a membrane permeable ion exchanger, exchanging luminal protons for cations (Zhang et al., 1993). This rapid increase in osmotically active cations leads to observable TGN/EE swelling. The cation concentration gradient is likely to be an important factor determining the severity of monensin induced TGN/EE swelling, as the *nhx5/nhx6* mutants are less sensitive to monensin, despite a more acidic luminal pH and therefore, likely to have a greater proton gradient (Dragwidge et al., 2019). We hypothesised that if CCC1 is important for ion efflux out of the TGN/EE, *ccc1* TGN/EE might be hypersensitive to monensin-induced TGN/EE swelling. We assessed the susceptibility of *ccc1* to monensin using live-cell imaging of VHAa1-RFP-labelled TGN/EE in wildtype and *ccc1* root epidermal cells in the elongation zone, treated with 2.5 μM of monensin for 15 min. The average wildtype TGN/EE size increased by 28±13% while *ccc1* TGN/EE size increased by 52±16%, revealing a highly increased susceptibility of *ccc1* TGN/EE to osmotically induced swelling caused by cation accumulation (Fig. 7A). This suggests that CCC1 mitigates the effect of monensin in wildtype cells, probably by effluxing K^+^ and Cl^−^, as anion and cation transport are stochiometrically linked in CCC1. We further suspected that TGN/EE pH could be altered as a consequence of altered cation and anion transport processes.

**Figure 7.**
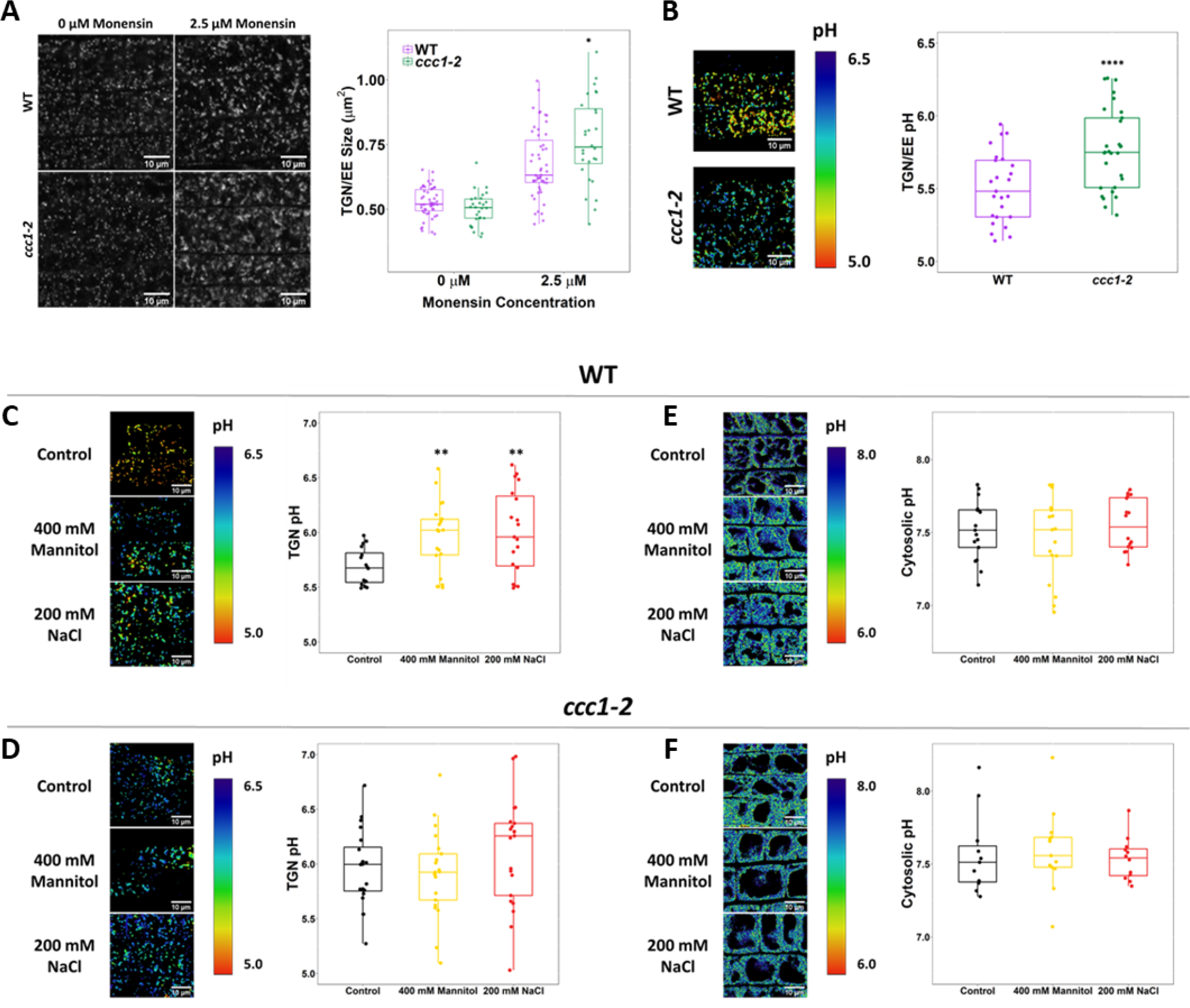
CCC1 is required for regulation of TGN/EE luminal conditions and luminal pH change in response to osmotic shock. A) Osmotically induced TGN/EE swelling due to cation influx is more severe in *ccc1* epidermal root cells aft er 15 min of treatment with 2.5 μM monensin. TGN/EE are visualised through stable expression of VHAal-RFP (white) in wildtype and *ccc1* backgrounds. n > 28 plants. Scale bars are 10 μM. Images are single representative optical sections. B-D) Stable expression of the TGN/EE lumen pH sensor pHusion in wildtype and *ccc1* backgrounds. C) The pH of *ccc1* TGN/EE is 0.3 pH units higher compared to wildtype in root epidermal cells. n > 26. D) The pH of Wt TGN/EE increased by 0.3 in response to 15 minute treatments with 400 mM mannitol and 200 mM NaCl. n > 16. E) The same treatments resulted in no change to the pH of *ccc1* TGN/EE. n > 16. E-F) Stable expression of the cytosolic pH sensor pHGFP in wildtype and *ccc1* backgrounds. There is no significant pH difference between the two genotypes in root epidermal cells. The cytosolic pH or both E) Wt, and F) *ccc1* was unchanged in response to osmotic and salt shock. n > 13. pH with both sensors was calculated using calibration curves performed with each set of experiments. Images show the ratio of GFP/RFP of the pH-sensor SYP61-p Husion in the TGN/EE lumen and emission ratio for the pHGFP cytosolic sensor as a colour range, using the inverted “physics” look-up table in lmageJ. Scale bars are 10 μM. Boxplots show range excluding outliers; median, 1st and 3rd quartile are indicated. Points represent individual measurements. Student’s t-tests comparing *ccc1* to wildtype. * indicates P < 0.05, ** indicates P < 0.01, **** indicates P < 0.0001.

**Figure 8.**
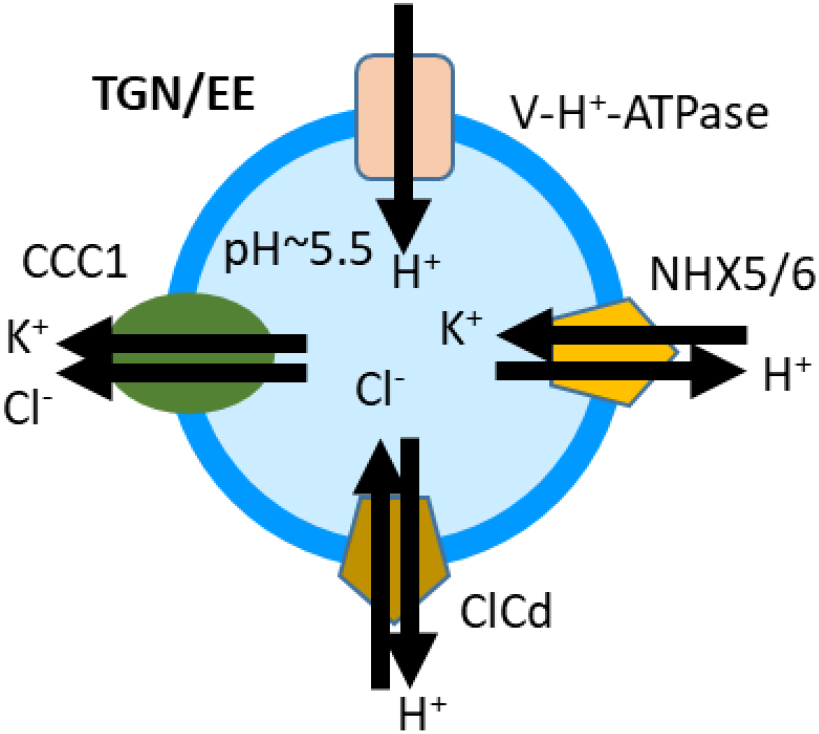
Prnposed model of ion and pHI iregulation in the TGN/EE. The V-H^+^-ATPase proton pump, the cation proton exchangers NHXS and NHX6, and the anion proton exchanger CLCd have been previously shown or proposed to be important for pH regulation in the TGN/EE lumen. CCC1 is the first candidate for providing a cation and anion efflux mechanism, completing the regulatory transport circuit.

To investigate this we introduced the stably expressed TGN/EE localised pH-sensor, SYP61-pHusion (Luo et al., 2015), into *ccc1* and investigated if defects in cation anion symport can impact pH regulation in the TGN/EE. Confocal imaging of epidermal cells in the root elongation zone revealed that luminal pH was more alkaline in *ccc1,* with a pH of 5.8 ±0.05, compared to wildtype pH of 5.5 ±0.05, confirming our initial hypothesis that a non-proton transporting ion symporter can impact luminal pH (Fig. 7B). We then investigated if loss of CCC1 might lead to a more general defect of intracellular pH. Therefore, we measured vacuolar pH using the pH sensitive dye, BCECF (Krebs et al., 2010). No difference was found between the genotypes, demonstrating that lack of CCC1 leads to spatially defined pH changes instead of a general effect (Fig. 7E-F, S4). To further assay the role of CCC1 in the TGN/EE and how it may interact with other TGN/EE transporters and the proton pump, crosses were made of *ccc1* × *det3* and *ccc1* × *nhx5 × nhx6*. However both crosses resulted in plants which were not viable. This might further support that CCC1 is required for the regulation of the TGN/EE luminal conditions in concert with other ion and proton transporters, and that removal of more regulatory components leads to such severe defects that the plants are unable to survive.

### The luminal pH of the TGN/EE increases in response to salt and osmotic stress

Plants with altered TGN/EE lumen pH show alterations in response to salt and osmotic treatments, suggesting that this regulation is important in wildtype plants in response to these stresses. We therefore investigated first in wildtype if there are changes to TGN/EE luminal pH in response to salt and osmotic treatment, and then compared our findings with *ccc1* knockouts.

The TGN/EE luminal pH of epidermal root cells of the elongation zone was quantified after 15 min exposure to 200 mM NaCl or 400 mM mannitol, which was compared to the pH of plants exposed to just the 1/2 MS control solution. In wildtype, in response to both NaCl and mannitol, the pH of the TGN/EE increased from 5.7 ± 0.04 (SEM) under control conditions to 6.0 ± 0.09 and 6.0 ± 0.07 respectively (Fig. 7C). Mannitol and NaCl both elicited the same pH response from plants demonstrating that the pH change is not exclusively connected to an increase in external ion concentrations but rather, occurs under an elevated external osmolality as well. Osmotic stress triggers an array of responses at the PM that can result in membrane depolarisation and depletion of cellular ATP (Che-Othman et al., 2017). Such events may impact proton transport across multiple membranes and as such, pH changes under osmotic stress may not be exclusive to the TGN/EE lumen. To assess this, the pH of the cytosol was also measured in response to stress. We used the genetically encoded and stably expressed pHGFP (Moseyko et al. 2001) and excluded the signal from nuclei for measurements, because the pH of the nucleus differs from that of the cytosol and may therefore also differ in changes in response to stress (Shen et al., 2013). We found that neither 15 min NaCl nor mannitol treatment induced significant changes in cytosolic pH (Fig. 7E). Under control conditions, the pH of the wildtype cytosol was 7.5 ± 0.05. Almost identical pH values of 7.5 ± 0.07 under NaCl treatment and 7.6 ± 0.04 under mannitol treatment were obtained. In the same set of experiments, TGN/EE and cytosolic pH of *ccc1* was assayed.

As in wildtype, no changes were observed for cytosolic pH in *ccc1* (Fig. 7F). *ccc1* cytosolic pH under control conditions was 7.5 ± 0.12. When treated with NaCl, the pH was 7.6 ± 0.08 and when treated with mannitol the cytosol pH was 7.5 ± 0.04. In addition, cytosolic pH in wildtype and *ccc1* under control conditions are almost identical, further supporting the idea that the pH change in *ccc1* is compartment specific and not a generalised effect. Also in this set of experiments, the TGN/EE luminal pH of *ccc1* TGN/EE was higher than wildtype with a pH of 6.0 ± 0.08 in control conditions. When treated with NaCl or mannitol, no significant change in luminal pH was detected with mean TGN/EE pH values of 6.1 ± 0.11 (SEM) and 5.9 ± 0.08 in response to NaCl and mannitol, respectively (Fig. 7D).

Collectively, these results suggest that there is a TGN/EE specific pH response to osmotic stress in wildtype plants, whereas no change in TGN/EE pH was found in *ccc1*, which might indicate that the pH is set to 6.0 in response to stress or that pH 6 may be an upper limit for the TGN/EE.

## Discussion

In the TGN/EE, both an increase or a decrease of luminal pH leads to changes in endomembrane trafficking, cell expansion, and cell wall formation, which implies that the pH of the TGN/EE is finely regulated. Here, we show that CCC1 is localised in the TGN/EE in Arabidopsis, with no evidence found of PM localisation; and that loss of the cation anion symporter leads to defects in establishing the typical, low pH in TGN/EE lumen. This makes CCC1 the first TGN/EE-localised transporter important for pH regulation that does not directly transport protons. Our results show that CCC1 is important for counteracting the cation-induced osmotic swelling of TGN/EE by monensin (Fig. 7), consistent with the symporter being important for regulating luminal ion concentrations. pH measurements confirmed that *ccc1* cells have a more alkaline pH in the TGN/EE lumen compared to wildtype. We therefore propose to extend the current model of pH regulation, containing the V-H^+^-ATPase, NHX5 and NHX6, and CLCd (Sze et al., 2018) with CCC1 (Fig. 6).

CCC1 is an electroneutral anion cation symporter, which fixes the stoichiometry of exported anions and cations at 1:1 (Colmenero‐Flores et al., 2007). This ion export ratio would therefore tightly connect the activities of the ion exchangers, and contribute to the strict regulation of pH. In addition, other transporters such as the potassium efflux antiporters KEA4, KEA5 and KEA6 might also contribute to luminal pH regulation *in planta*; yet their roles are currently less understood since triple *kea4/kea5/kea5* knockouts show a much less severe phenotype compared to *nhx5/nhx6*, *det3*, or *ccc1* (Dettmer et al., 2006, Colmenero‐Flores et al., 2007, Bassil et al., 2011, Zhu et al., 2018). Recent work suggests that CLCd and CLCf have partially redundant roles and effect trans-Golgi pH instead of TGN/EE pH where they are localised, demonstrating the complexity of the regulatory network (Scholl et al., 2021)

Functional endomembrane transport has a vital role in many cellular processes, including cell wall formation, nutrient acquisition and the establishment of hormone gradients (Guo et al., 2014, Adamowski and Friml, 2015, Sinclair et al., 2018). In agreement with this, *ccc1* mutant plants display very severe phenotypic alterations compared to wildtype. We found that CCC1 is ubiquitously expressed, and its function is required for cell elongation, which might be the cause of the reduced overall growth of the mutants (Fig. 2, S1). Cell elongation is also reduced in *nhx5/nhx6* and *clcd/amiR-clcf* mutants; with reduced root hair elongation speed in both and reduced root epidermal cell and root hair length in *clcd/amiR-clcf* (Bassil et al. 2011; Scholl et al., 2021). Reduced root hair elongation could be the result of reduced rates of endomembrane trafficking, which have been observed in *nhx5/6* causing reduced delivery of cargo important for cell expansion such as cell wall components. Phenotypic defects in *ccc1* are not restricted to cell elongation, we found that trichoblast patterning was also altered in *ccc1* roots (Fig. S1). *ccc1* plants further exhibit highly increased shoot branching, which gives the knockout plants a bushy appearance (Henderson et al., 2018). One cause for this might be defects in auxin transport regulation in shoots. We found alterations in PIN2 exocytosis in *ccc1* roots, suggesting that similar defects could exist in shoot cells (Fig. 5).

Previous work has revealed a link between endomembrane trafficking, the TGN/EE and salt tolerance in plants. Knockouts of the Arabidopsis Rab GTPase ARA6, involved in TGN/EE to PM trafficking, results in salt sensitive plants. Likewise, knockouts of VACUOLAR PROTEIN SORTING 4 (VPS4), involved in sorting of proteins for vacuolar degradation, results in salt sensitive plants. Similarly, both *det3* plants and RNAi knockdowns of the TGN/EE V-H^+^-ATPase, display a stronger reduction in primary root growth in response to 50 mM NaCl compared with wildtype plants (Batelli et al., 2007; Krebs et al., 2010). *nhx5/nhx6* plants exhibit severe reductions in growth when transferred to media containing 100 mM NaCl and germination of *nhx5/nhx6* seeds, with a reduced TGN/EE luminal pH, is almost completely abolished on media containing 100 mM NaCl (Bassil et al., 2011).

CCC1 appears to play a contrasting role to NHX5/NHX6, as here, we found that the germination of *ccc1* seeds was more tolarant to salt and osmotic strss than wildtype (Fig. 4). The contrasting roles in osmotic tolerance may be explained by the contrasting roles of the proteins in the TGN/EE. *nhx5/nhx6* knockouts have a lower TGN/EE pH and are more resistant to monensin, while *ccc1* knockouts have a higher TGN/EE pH and are hypersensitive to monensin. NHX5 and NHX6 import K^+^ from the cytosol and therefore knockouts are likely to have a reduced accumulation of K^+^ in the TGN/EE wheras we find here that CCC1 is likely to export K^+^ and therefore knockouts likely have an increased accumulation of K^+^.

The root hairs of *ccc1* plants elongate slower than wildtype under control conditions, however, when challenged with an elevated osmolality, *ccc1* root hairs elongate at the same rate as wildtype root hairs. This result may suggest that CCC1 is important for root hair growth under standard conditions but less important at an elevated osmolality for cell elongation rate.

The greater germination rate of *ccc1* seeds on media with a higher osmolality may be the result of weaker cell wall and potentially a weaker seed coat, which might require a smaller water uptake for the necessary seed swelling and seed coat rupture to occur. Changes in the cell wall composition of *ccc1* knockouts has been observed (Han et al. 2020). Alternatively, improved tolerance of seed germination to an external osmolality could be related to abscisic acid (ABA) insensitivity. ABA insensitivity results in reduced osmotic stress induced inhibition of germination and the endomembrane system can impact ABA sensitivity through the delivery of PM ascociated ABA receptors (Yuan et al., 2011; Belda-Palazon et al., 2016). Interestingly, overexpression of Arabidopsis NHX5 in both soybean (*Glycine max*) and paper mulberry (*Broussonetia papyrifera* L. Vent) results in improved salt tolerance of plants and enhanced proline accumulation. Overexpression of NHX5 in paper mulberry also resulted in improved drought tolerance and an increased accumulation of Na^+^ and K^+^ in leaves (Li et al., 2011; Wu et al., 2016).

Knockouts of *ccc1* require higher concentrations to induce plasmolysis of root epidermal cells. This increased tolerance to plasmolysis could occur if *ccc1* knockouts were able to regulate their internal osmolality faster than wildtype. It is possible that *ccc1* root epidermal cells could influx external osmolytes faster, due to altered localisation of osmolyte transporters, to more rapidly match the external osmolality. Supporting this, nutrient uptake and ion translocation have been shown to be affected in *ccc1* plants, with altered total shoot K^+^, Na^+^ and Cl^−^ accumulation compared to the wildtype (Colmenero‐Flores et al., 2007; Henderson et al., 2015). Interestingly, *Osccc1.1* roots have a lower cell sap osmolality, but an increase in biomass was reported when plants grown in saline conditions compared to control conditions (Chen et al. 2016). The model we propose for Arabidopsis CCC1 might therefore be similar in rice, suggesting a conserved function.

In additionally, overexpression of trifolate orange CCC, *PtcCCC* in tobacco plants results in plants with improved shoot biomass in response to 200 mM KCl and improved root biomass in response to 200 mM KCl or NaCl (Wei et al., 2018). Furthermore, in a screen of the seed ionome, *ccc1* seeds were identified as containing high concentrations of Fe, Ca and S, and low K and Na (McDowell et al., 2013). The alterations in Fe, Ca and S are of interest as these are not substrates of CCC1, which suggests loss of CCC1 function impacts the capacity of PM and/or tonoplast transport due to defects of TGN/EE function, most likely through impacts on trafficking.

Here we show the TGN/EE pH of epidermal root cells increases in response to both salt and osmotic shock, which has not been reported previously. Endomembrane trafficking and the abundance of PM proteins such as PIN1, PIN2 and BRI1 is altered when the pH of the TGN/EE is changed (Luo et al., 2015; Reguera et al., 2015; Dragwidge et al., 2018; Dragwidge et al., 2019). Therefore, modulation of the TGN/EE pH in response to abiotic stress may enable plants to rapidly regulate protein trafficking and abundance. *det3*, *nhx5/nhx6* and *ccc1* plants all exhibit TGN/EE pH differences to the wildtype of around 0.3 units, very similar to what we consistently found for *ccc1*.

Our research here provides evidence for a central role for CCC1 in cellular function, and fills a gap in our understanding of how endomembrane luminal pH is regulated through the identification of a novel ion efflux component of the TGN/EE membrane transport network. This regulation is dynamic and there is a specific pH response to osmotic and salt shock, which is disrupted in CCC1 loss of function plants.

## Material and Methods

### Plant material and growth conditions

*Arabidopsis thaliana* plants were all in the Columbia-0 (Col-0) background. Previously described T-DNA insertion lines in AT1G30450, *ccc1-1* (SALK-048175), and *ccc1-2* (SALK-145300) were used in this study (Colmenero‐Flores et al., 2007). *PIN2::PIN2-GFP, 35S::VHAa1-RFP*, *UBQ10::PIP2;1-GFP, 35S::GFP-LTIB6 and 35s::SYP61-pHusion* constructs and plant lines were previously described (Cutler et al., 2000; Xu and Scheres, 2005; Dettmer et al., 2006; Luo et al., 2015).

Arabidopsis plants were grown on half strength Murashige and Skoog (1/2 MS) media containing 0.1% sucrose, 0.6% phytagel, pH 5.6 with KOH. Plants were sown on plates, incubated at 4°C for at least 2 d and subsequently grown vertically at 21°C and 19°C in 16 h light and 8 h dark, respectively; or 8 h light and 16 h dark when short day is indicated. Plants were grown for different periods of time, as indicated below and in figure legends. Plants in pots (Fig. 2), were grown in Irish Peat soil.

### Promoter activity analysis by GUS and Venus fluorescence

GUS staining was done according to Jefferson et al. (1987). In summary, plants were submerged in GUS-staining solution and stained (plant age and staining times indicated in figure legend). Image of the entire rosette was captured with a Nikon digital camera, flower and inflorescence images with a Nikon SMZ25 stereo microscope. Fluorescence of the nuclear localised NLS-Venus was imaged in plants ranging from 5–8 d to 8 weeks as indicated in figure legend. Excitation light wavelength was 514 nm, emission was detected at 520–560 nm, using either a Nikon A1R or an Olympus FV3000 Confocal Laser-Scanning Microscope; with the following objectives: 20× Plan Apo Lambda and 40× Apo LWD WI Lambda S (Nikon), and 10× UPSLAPO objective (Olympus).

### Root hair length, root hair elongation rate

Light microscopy imaging of root hair length was performed using a Nikon SMZ25 stereo microscope with a 2× objective. For quantification of root hair length, images of roots were taken from above the maturation zone of 6 d plants. Each measurement was of a single root hair. Multiple root hairs were measured per plant. Root hair length was measured using FIJI (Schindelin et al., 2012). For time-lapse light microscopy of root hair elongation rate, plants were germinated in 2 mL of media placed in 1-well microscopy slides (Thermo Fisher) and grown vertically. Images of root hairs in the maturation zone were taken every 30 s for 6 h using a Nikon Diaphot 300. Measurements were taken from the beginning of root hair elongation, of root hairs that elongated beyond the initiation phase, until root hair growth ceased. For consistency, elongation rates of root hairs were only measured for root hairs where both initiation and cessation of growth could be observed in the time-lapse. A single root hair was measured per plant. Analysis and creation of videos was performed using FIJI.

### Root morphology imaging

Root morphology images for epidermal cell length and ectopic root hairs were taken at the same Nikon confocal, using 6 d seedlings and root cell wall autofluorescence (excitation 404 nm, emission 425–475 nm). Each measurement for epidermal cell length was of a single cell. Multiple cells were measured per plant.

### GFP-CCC1 cloning and expression

For stable expression of *CCC1* in root hairs, 1402 bp of the trichoblast-specific promoter *EXP7* (Marquès‐Bueno et al., 2016) was first amplified from Col-0 genomic DNA, using the primers EXP7pro-HindIII_F (*tatacAAGCTTATTACAAAGGGGAAATTTAGGT*) and EXP7pro-KpnI_R (*cttatGGTACCTCTAGCCTCTTTTTCTTTATTC*), following a Phusion^®^ PCR protocol (NEB). The PCR product and the binary plasmid pMDC43 were subsequently cut with the restriction enzymes *Hind*III-HF and *Kpn*I-HF to remove the *2×35S* promoter, and the digestion reactions were purified using illustra^TM^ GFX^TM^ PCR DNA and Gel Band Purification kits. Fragment ligation was performed using T4 DNA Ligase protocol (NEB) at 16°C overnight. 2 μl of the ligation reaction was transformed into *E. coli* DB3.1 cells and after a sequencing verification, a plasmid was subsequently selected that showed the correct replacement of the *2×35S* promoter with the *EXP7* promoter. *CCC1* CDS (with stop codon) was then shuttled into the pMDC43EXP7 using LR clonase II enzyme, which creates N-terminally *GFP*-tagged *CCC1*. Correct plasmids were transformed into *Agrobacterium tumefaciens*, and heterozygous Arabidopsis plants (*ccc1*^+/-^) were floral dipped as the homozygous *ccc1* knockout does not support floral dipping well. Floral dipping was performed according to Clough and Bent (1998), and transformants were selected on 1/2 MS plates with no sucrose, containing hygromycin for selection. Homozygous *ccc1* knockouts expressing *GFP-CCC1* were selected, and phenotyped in the next generation.

### Colocalisation

For colocalisation, 6 d plants expressing both *GFP-CCC1* (excitation 488 nm, emission 500– 550 nm) and the TGN/EE marker VHAa1-RFP (excitation 561 nm, emission 570–620 nm) were imaged using Nikon A1R Confocal Laser-Scanning microscope, using a 60× Plan Apo VC WI objective with a numerical aperture of 1.2; pinhole set to of 1.2 AU. Five roots were imaged, with three images of three separate mature epidermal root cells being imaged per plant. Analysis was performed on image stacks with a step size of 0.45 μm. Colocalisation was assessed using the FIJI plugin DiAna (Gilles et al., 2017). In brief, segmentation was performed using the iterative segmentation function before the number of objects that overlap are counted. The percentage of overlapping objects is reported as the percentage of colocalisation. Colocalisation with DiAna was supported by obtaining the Pearson’s coefficient on the same stacks using JACoP (Bolte and Cordelieres, 2006).

### Germination and collet hairs

Both seed germination and collet hair formation was assayed on the same set of seeds. Seeds were germinated on 1/2 MS media containing the indicated concentration of mannitol, sorbitol or NaCl. The control seeds, germinated on media with no added mannitol, sorbitol or NaCl, were shared between all three treatments. Plants were stratified at 4°C for 2 d, and then transferred to a growth chamber. 6 d after transfer, the percentage of seeds that had germinated were counted with radicle emergence being used as the indicator for germination. On the same day, for all plants that had developed to the point where collet hairs should be visible, the percentage with properly developed collet hairs was determined. Development of collet hairs was scored based on 4 categories, plants with no visible collet hairs, plants with one or two collet hairs present, plants with a cluster of collet hairs but lacking a complete ring of collet hairs and plants with a complete ring of collet hairs.

### Root cell sap osmolality

For each sample, seeds (approx. 50 per plate) were placed in a row at the top of a 1/2 MS media plate. After 12 d, roots were cut from the shoots and all roots from the same plate placed together in tubes with 100 μL of water. The sealed tubes were heated to 85°Cfor 20 min. Tubes were cooled before osmolality was measured with a Friske 210 micro-sample osmometer. Final displayed osmolality was adjusted for the fresh weight of the input material.

### Root cell sap osmolality

For each sample, seeds (approx. 50 per plate) were placed in a row at the top of a 1/2 MS media plate. After 12 d, roots were cut from the shoots and all roots from the same plate placed together in tubes with 100 μL of water. The sealed tubes were heated to 85°C for 20 min. Tubes were cooled before osmolality was measured with a Friske 210 micro-sample osmometer. Final displayed osmolalitywas adjusted for the fresh weight of the input material.

### Plasmolysis

Mannitol was used to induce plasmolysis of plants grown for 6 d on 1/2 MS without mannitol. The mannitol concentration at which root cells plasmolysed was determined using epidermal cell without root hairs, in plants expressing the PM marker GFP-LTI6b. Plants were transferred from the growth media to a liquid 1/2 MS solution containing different concentrations of mannitol, as indicated in Fig. 3, 1 h before counting. Plasmolysed and non-plasmolysed cells were counted under a Nikon Ni-E widefield microscope. From each root, the plasmolysis state of 40 cells was assessed. Cells which had root hairs were not included. A cell was considered as plasmolysed if the corners of the cell had detached/curved. Only cells past the maturation zone of the plant were included. The estimated concentration at which 50% of cells would plasmolyse for each genotype was determined by fitting a Boltzmann sigmoid curve in Graphpad Prism 9 to obtain the ‘V50’.

### Trafficking of PIP2;1-GFP

Internalisation of PIP2;1-GFP in response to osmotic stress was measured in epidermal root cells of the elongation zone of 6 d old plants. Plants were transferred from 1/2 MS media to liquid 1/2 MS containing 100 mM NaCl or 200 mM mannitol 30 min before imaging, or control 1/2 MS. Imaging was performed using a Nikon A1R confocal laser scanning microscope with a 60× Plan Apo VC WI objective. A ratio of PIN2;1-GFP (excitation = 488 nm, emission = 500 – 550 nm) at the PM and inside the cell was determined by measuring the signal intensity at the PM and inside the cell with FIJI. All results were normalised to wildtype median signal ratio under control conditions.

### Endo- and Exocytosis

Endocytosis was assayed in the root tips of 6 d plants using the fluorescent membrane stain FM4-64 (excitation 561 nm, emission 570–620 nm) and the endomembrane trafficking inhibitor brefeldin A (BFA). Plants were either incubated in 1/2 MS containing 4 μM FM4-64 and 25 μM BFA in the dark for 10 min before imaging or for 10 min in 1/2 MS with 4 μM FM4-64 and 25 μM BFA in the dark before washing and incubating in 1/2 MS with 25 μM BFA for 50 min before imaging. Images were single representative optical sections, taken of epidermal cells in the root elongation zone. A ratio of cytoplasm:PM signal was measured in ImageJ by using the polygon selection tool to measure the mean grey value of the entire interior of the cell and divide this by the PM mean grey value, acquired using the segmented line tool (width 1).

Trafficking of FM4-64 to the tonoplast was measured by incubating plants in 1/2 MS containing 4 μM FM4-64 for 10 min before washing and incubating in 1/2 MS for 3 h. Images were single representative optical sections taken of epidermal cells in the root elongation zone. A ratio of tonoplast:PM signal was acquired using the segmented line tool for both PM and tonoplast measurement.

Exocytosis was assayed in the root tips of 6 d old plants using PIN2-GFP or LTI6b-GFP in the epidermis. For the “t0” image point, plants were taken directly off growth media and immediately imaged. Otherwise, plants were treated with 25 μM BFA for 60 min, at which point some plants were imaged to obtain the “60’ BFA” images. The rest were washed in liquid 1/2 MS and left to recover for 60 min in liquid 1/2 MS media before imaging for the “60’ washout” images. Signal internalisation was measured as described for endocytosis. Imaging was done with the 60× objective and Nikon A1R Confocal Laser-Scanning microscope described above.

### TGN/EE swelling with monensin

TGN/EE swelling was induced using the ionophore, monensin (Sigma). 6 d plant roots expressing VHAa1-RFP (excitation 561 nm, emission 570–620 nm) were submerged in ½ MS solution with or without 2.5 μM monensin for 15 min before imaging. Imaging was performed on the Nikon A1R Confocal Laser-Scanning microscope and 60× objective described above. Images are single representative optical sections, taken of epidermal cells in the root elongation zone. A single measurement was taken per plant, which included multiple cells. Measuring was done using FIJI. TGN/EE were identified using VHAa1-RFP signal and segmented using the automatic threshold algorithm, “RenyiEntropy”. The average size of the segmented TGN/EE was measured using “Analyse Particles”.

### TGN/EE and cytosolic pH measurements

The pH of the TGN/EE was measured as described in Luo et al. (2015). In brief, SYP61-pHusion (excitation 488 and 561 nm, emission 500–550 and 570–620 nm) was imaged in epidermal cells of the root elongation zone of 6 d plants on the Nikon A1R Confocal Laser-Scanning microscope and 60× objective described above, obtaining a single optical section. GFP/RFP intensity ratios were obtained from cells incubated in solutions of known pH to create a calibration curve. TGN/EE pH was measured by obtaining ratios from control plants. The calibration curve was created by measuring cells treated for 15 min with 50 mM MES-BTP (bis-tris-propane) or 50 mM HEPES-BTP with 50 mM ammonium acetate. Seven points between pH 5.0 and 8.0 were measured for the calibration curve. Six measurements were taken per point for calibration and a calibration was performed before every experiment. The curve was fit using a Boltzmann sigmoidal in GraphPad Prism 8.0. A single measurement was taken per plant, which included multiple cells. The GFP/RFP ratio was obtained in FIJI by firstly segmenting for TGN/EE using the automatic “RenyiEntropy” threshold on the RFP signal before measuring fluorescent intensity.

The cytosol was measured as described above for the measurement of the TGN/EE with some adjustments. The cytosolic pH sensor pHGFP (excitation 405 and 488 nm, emission 500–550 nm; Moseyko et al. 2001) was used. Sections of images used for measurement were manually cropped to remove any nuclei. pH measurements were obtained by acquiring a ratio of 488/405 nm signal intensity. Automatic thresholding in FIJI was performed using the “default” algorithm on the 488 nm channel.

For TGN/EE and cytosolic pH measurements after treatment with osmotic or salt shock, plants were submerged in 1/2 MS containing 400 mM mannitol or 200 mM NaCl for 15 min before imaging.

### Cell types

Imaging for pH and monensin measurements, as well as for endo- and exocytosis, was done in epidermal cells in the root elongation zone; imaging of GFP-CCC1 for colocalisation was done in mature epidermis cells, specifically in trichoblasts.

## Acknowledgments

We thank Melanie Krebs and Karin Schumacher for providing seeds expressing the TGN/EE and cytosolic pH sensors, and Chuang Wang for the construction of PIP2;1-GFP seeds. Matthew Tucker for the 3×VenusNLS plasmid, Steve Tyerman for helpful discussions, Philip Brewer for *PIN2::PIN2-GFP* seeds, and Christian Luschnig for advice on genotyping. We thank Renée Phillips and Marie Beillevert for assistance with lab and plant work. We thank Adelaide Microscopy, especially Gwen Mayo and Jane Sibbons, for support with microscopy; and we thank the University of Melbourne Advanced Microscopy Facility where electron microscopy was conducted. We thank the Australian Research Council for funding this work through DE170100054 to HEM, FT130100709 and CE140100008 to MG, and DE160100804 to SW; HEM is also supported in part by funding from the CRC program as the Canada Research Chair in Plant Cell Biology.

## Author contributions

SW led the project; DWM, MG and SW designed experiments; DWM conducted most experiments with contributions from SW, HEM, YQ and AS; DWM, MG and SW wrote the paper, HEM, AS and YQ commented on the paper.

## Supplementary Materials and Methods

### Root morphology imaging

Root morphology images for epidermis cell length and root hair cell files were taken at a Nikon A1R Confocal Laser-Scanning microscope, using 6 d old seedlings and root cell wall autoflourescence (excitation = 404 nm, emission = 425-475 nm). Adjacent trichoblasts were also imaged in root epidermal cells of the maturation zone of *ccc1* plants expressing the trichoblast marker *PRP3::H2B-2xmCherry* (excitation = 561 nm, emission = 570-620 nm; Marques-Bueno et al. 2016).

### GFP-CCC1 imaging

6 d old plants expressing GFP-CCC1 (excitation = 488 nm, emission = 500-550 nm) driven by the *EXP7* promoter were imaged on a Nikon A1R Confocal Laser-Scanning microscope. For observation of GFP-CCC1 within BFA bodies, roots were first incubated in 1/2 MS with 4 μM FM4-64 (excitation = 561 nm, emission = 570-620 nm) for 5 min before being washed and incubated in 1/2 MS with 25 μM BFA for 1 h. Stacks were taken of elongating root hairs. For observing if GFP-CCC1 cycles to the PM, plants were incubated for 30 min in 1/2 MS with or without 50 μM tyrphostinA23. Stacks were taken of mature root epidermal cells. For observing if osmotic shock can induce a change of localisation of GFP-CCC1, plants were incubated in 1/2 MS solution for 1 h before being transferred to milliQ water, 1/2 MS solution or 1/2 MS solution containing 300 mM sorbitol for 5 min before imaging. Stacks were taken of elongating root hairs.

### High-pressure freezing and transmission electron microscopy (TEM)

Wild type and ccc1 mutant seeds were grown on 1/2 MS plates with 1% sucrose in long-day conditions for 5 d. Roots tips were excised using a sharp razor then cryofixed according to McFarlane et al. (2008), with modifications. Cryofixation was performed using a Leica EMICE high pressure freezer, type A carriers (Leica), and 1-hexadecene (Sigma) as a cryoprotectant. Roots were freeze substituted in a Leica AFS2 at −85 °C over 5 d in a mixture of 2% osmium tetroxide (Electron Microscopy Sciences) and 8% 2,2-dimethoxypropane (Sigma) in anhydrous acetone. Samples were warmed to room temperature over 2 d, then slowly infiltrated with Spurr’s resin over 5 d (Spurr, 1969). Samples were polymerized for 36 h at 70 °C, then sectioned to ~70 nm (silver) on a Leica UC7 microtome using a DiATOME knife. Sections were placed onto copper fine-bar grids (Gilder) coated with 0.3% formvar, then post-stained with 1% aqueous uranyl acetate (Polysciences) for 10 min and triple lead (sodium citrate, lead acetate, lead citrate from BDH, lead nitrate from Fisher; Sato, 1968) for four min. Samples were viewed using a Phillips CM120 BioTWIN transmission electron microscope with tungsten filament at an accelerating voltage of 120 kV and a Gatan MultiScan 791 CCD camera.

### Vacuole pH measurements

The pH of the vacuole was measured using BCECF-AM (Invitrogen) as described by Luo et al. (2015). In brief, 6 d old plants were incubated in the dark for 1 h in a 1/2 MS solution containing 10 μM BCECF and 0.02% pluronic F-127 (Invitrogen). Plants were washed before being imaged at the excitations 404 nm and 488 nm, both emissions were collected at 500-550 nm. The ratio between 404 and 488 is used to measure pH. A calibration curve was created by measuring cells treated for 15 min with 50 mM MES-BTP (bis-tris-propane) or 50 mM HEPES-BTP with 50 mM ammonium acetate. Seven points between pH 5.0 and 8.0 were measured for the calibration curve. Six measurements were taken per point for calibration and a calibration was performed before every experiment. The curve was fit using a Boltzmann sigmoidal in GraphPad Prism 8.0. A single measurement was taken per plant which included multiple cells. The 404/488 ratio was obtained in ImageJ by firstly segmenting using the automatic “default” threshold on the 488 nm channel before measuring fluorescent intensity.

---

**Figure S1.**
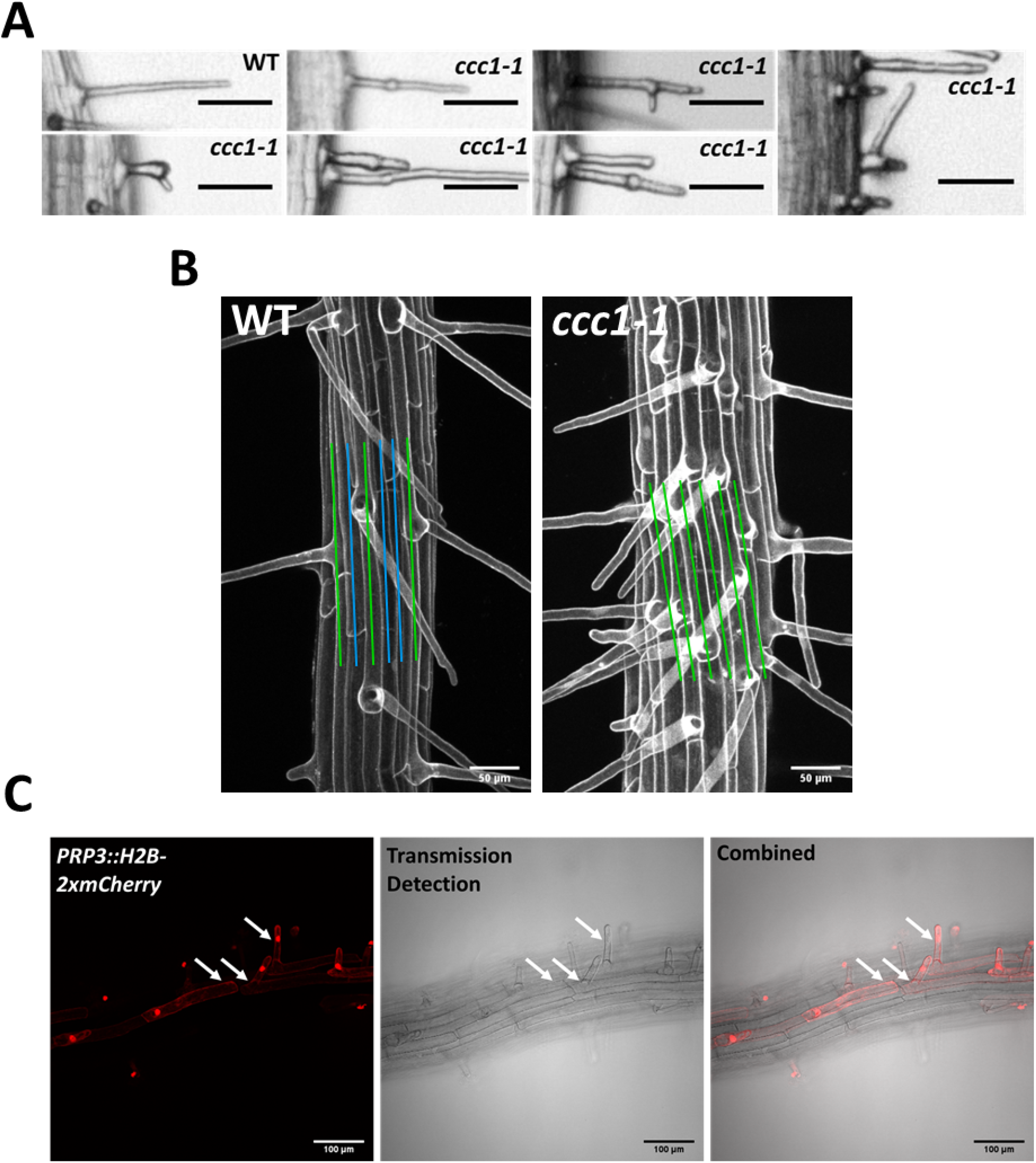
CCC1 is important for normal trichoblast development. A) A small number of *ccc1* root hairs display forking and branching. Scale bars are 50 μm. B) *ccc1* roots do not exhibit the classic cell file patterning of trichoblast and atrichoblast cell files observed in Arabidopsis. Cell files containing root hairs are marked with green while those that do not have root hairs are marked with blue. Images are of a maximum intensity projection, imaging cell wall autofluorescence. Scale bars are 50 μm. C) The trichoblast marker *PRP3::H2B-2xmCherry* (mCherry in red) confirms the appearance of adjacent trichoblasts in *ccc1.* Arrows point to epidermal root cells in *ccc1* which are of different files which are all expressing the trichoblast marker. Scale bars are 100 μm.

**Figure S2.**
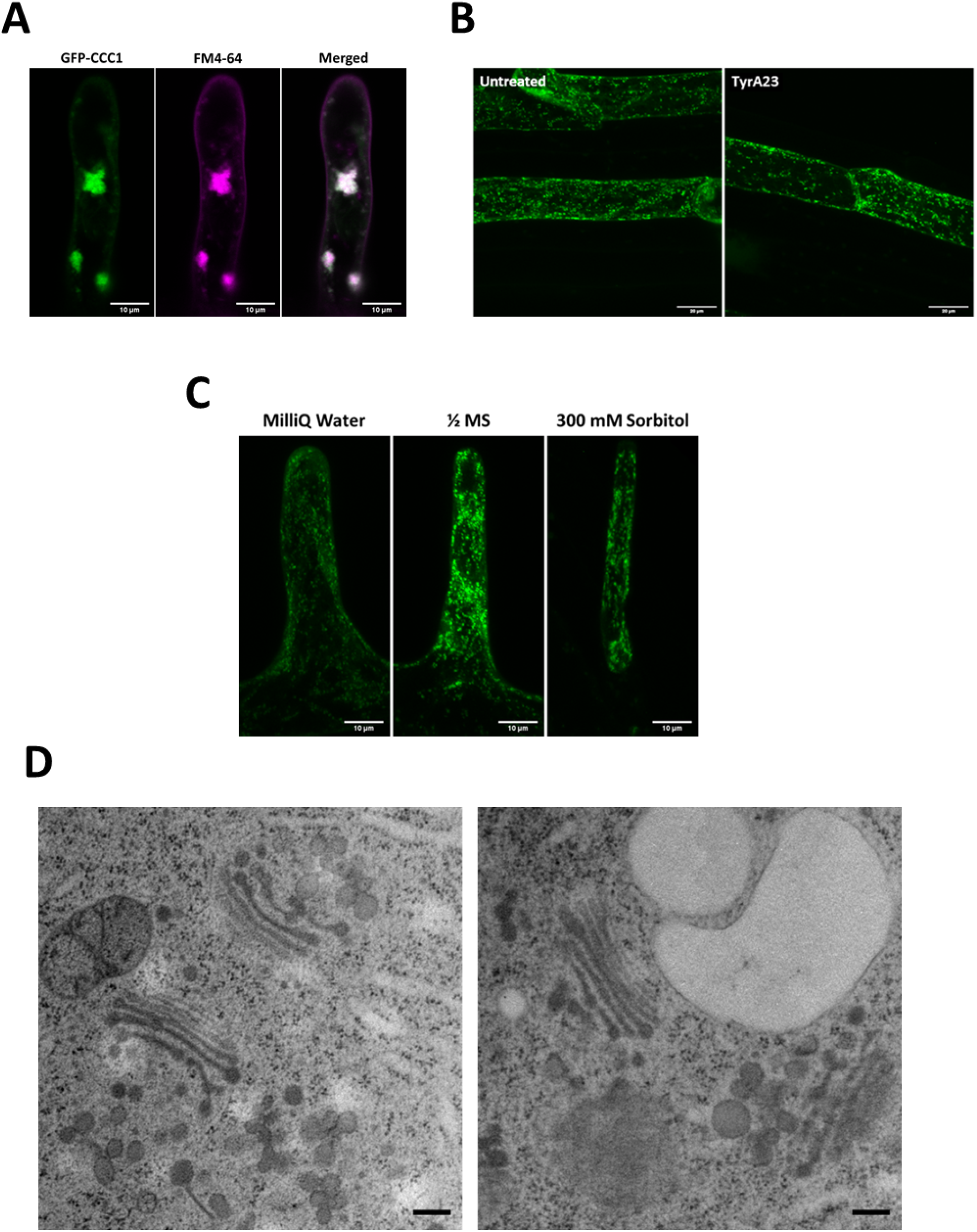
GFP-CCC1 is localised to the endomembrane system and does not cycle to the PM. A) After treating root hairs with 4 μM FM4-64 (magenta) for 5 min and 25 μM BFA for 1 h, GFP-CCC1 (green) and FM4-64 are both observed in the core of the BFA body. The image is a maximum intensity projection. Scale bars are 10 μm B) Treating roots with 50 μM of Tyrphostin A23(TyrA23) for 30 minutes, did not result in PM signal of GFP-CCC1 (green). Images are maximum intensity projections of mature root epidermal cells. Scale bars are 20 μm. C) Neither hyper- nor hypo-osmotic stress changed GFP-CCC1 localisation. Images are maximum intensity projections of root hairs equilibrated in 1/2 MS solution before being moved to MilliQ water or 300 mM sorbitol for 5 minutes. Scale bars are 10 μm. D) Representative TEM images of highpressure frozen, freeze substituted early elongation zone root epidermal cells, showing Golgi and TGN/EE morphology in Col-0 wildtype (left) and *ccc1* (right). Scale bars are 200 nm

**Figure S3.**
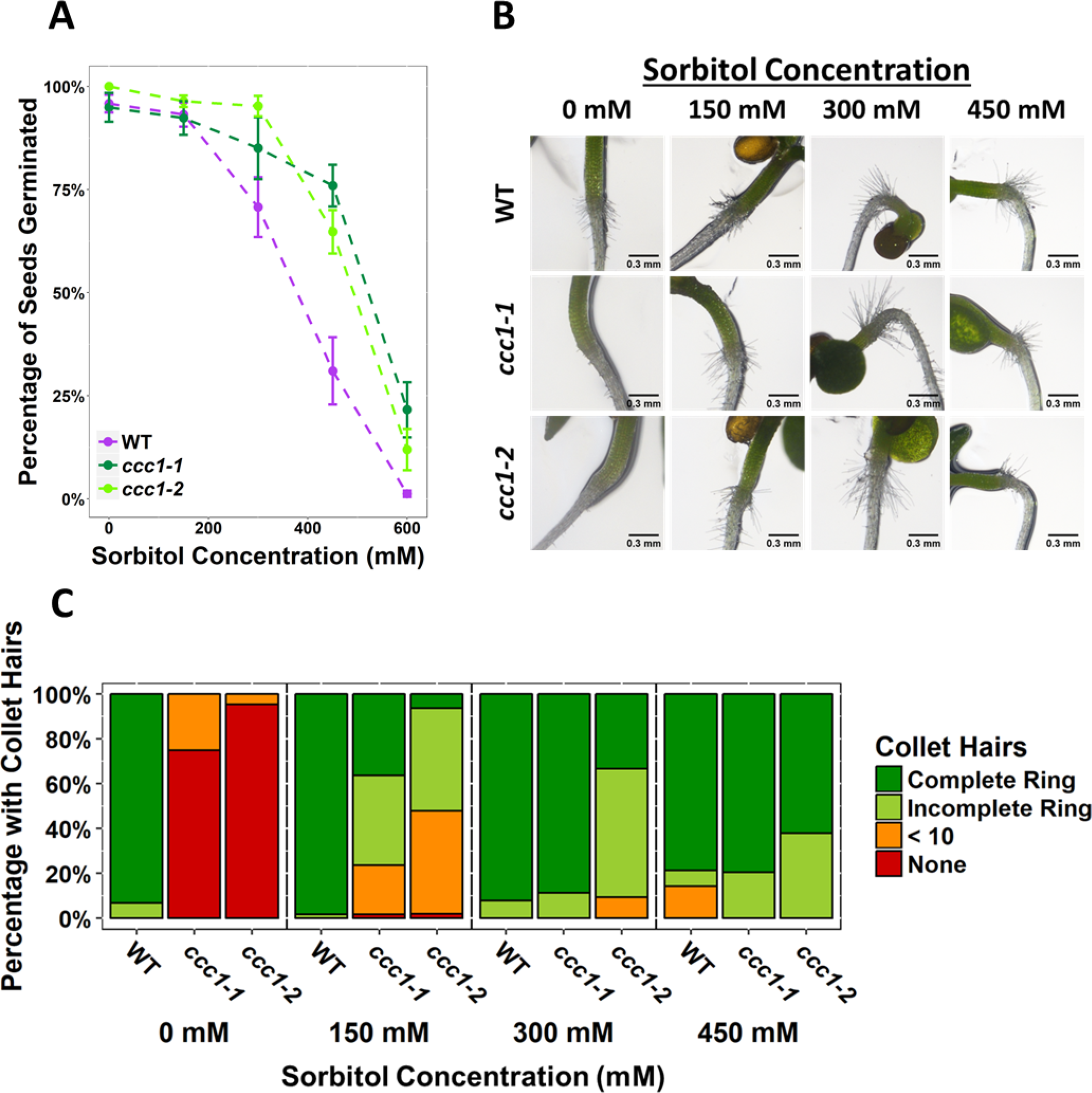
*ccc1* rnot hair elongation is rescued on sorbitol and germination of *ccc1* seeds is tolerant to osmotic stress imposed by sorbitol. A) A higher percentage of *ccc1* seeds germina te on media with a higher osmolality, adjusted with sorbitol. The number of germinated seeds assessed 6 d after imbibition. n > 90. B) Collet hair formation in *ccc1* is rescued on media with a higher osmolality. n > 9 plants for 450 mM treatments, > 30 plants for all other treatments. Scale bars are 300 μM. Germination and collet hair assays with mannitol, sorbitol and NaCl were performed together and therefore share the same control (0 mM).

**Figure S4.**
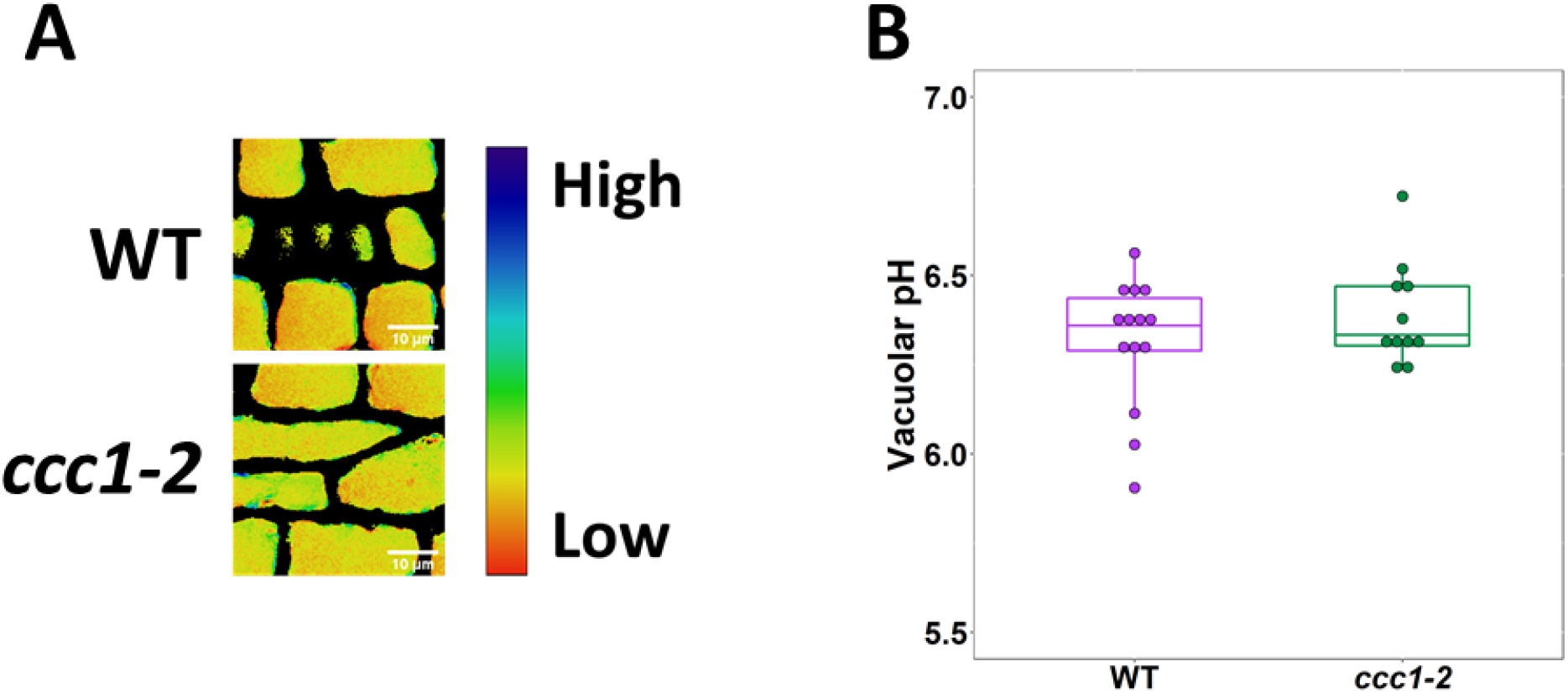
CCC1 does not regulate vacuolar pH. A) The pH of *ccc1* vacuoles is similar to wildtype in root epidermal cells, measured using the dye BCECF. Images show the ratio of the two BCECF excitations which were used to measure pH. n > 11 plants. Scale bars are 10 μM. Boxplot shows range excluding outliers; median and 1st and 3rd quartile are indicated. Points represent individual measurements. Students t-tests comparing *ccc1* to wildltype. * < 0.05

## Supplementary Video Legends

**Supplementary videos 1-2: Time lapse showing movement of GFP-CCC1 labelled subcellular compartments.** GFP-CCC1 localises to motile intracellular organelles in 3) root hairs and 4) trichoblasts, expression driven with EXP7 promoter. Time series of the root hair was taken through the centre plane of the root hair, while the trichoblast was imaged below the radial PM. Scale bars = 10 μm, 20 seconds shown per second at 10 frames per second.

**Supplementary videos 3-6: Time lapse of root hair elongation.** *ccc1* root hairs elongate slower than wildtype under control conditions and the same speed as wildtype when grown on media containing 150 mM mannitol. Scale bars are 100 μm, 48 min shown per second at 24 frames per second

